# The evolutionarily conserved APP-Spastin cooperation regulates endolysosomal homeostasis and apoptotic cell degradation

**DOI:** 10.64898/2026.07.14.738573

**Authors:** Qian Zheng, Fuqiao Liu, Lei Yuan, Zhulin Liu, Haibing Lv, Tong Xiao, Zheng Cui, Qiyuan Zhong, Hui Wang, Qiuyuan Yin, Hui Xiao

## Abstract

Efferocytosis, the recognition, engulfment, and degradation of apoptotic cells by phagocytes, is essential for tissue homeostasis and development, and its failure contributes to chronic inflammation and neurodegeneration. The amyloid precursor protein (APP), a central pathogenic factor in Alzheimer’s disease, retains physiological functions independent of amyloid production that remain poorly understood. Here, we identify a conserved, non-amyloidogenic role for APP in regulating apoptotic cell degradation via the endolysosomal pathway. Using *Drosophila* APPL as a model, structure-function analysis demonstrated that the intracellular internalization domain of APPL, but not its secreted ectodomain, is required for efficient apoptotic cell degradation. Immunoprecipitation coupled with mass spectrometry revealed a physical interaction between APPL and the microtubule severing ATPase Spastin, mediated by the microtubule-interacting and trafficking domain of Spastin. APPL interacts with Spastin on endosomal microtubules and modulates the dynamics of the Spastin-ESCRT-III complex, enabling Spastin to sever microtubules and promote endosomal tubule fission. Loss of APPL disrupts this process, causing aberrant endosomal tubulation and impaired lysosome biogenesis. Furthermore, it compromises the function of residual lysosomes, characterized by reduced acidity, diminished proteolytic activity, and increased lysosomal damage, which ultimately impairs the degradation of engulfed apoptotic cells. Critically, this phenotype is evolutionarily conserved in *C. elegans* and mice. Together, these findings establish a conserved APP-Spastin axis that regulates endolysosomal homeostasis and apoptotic cargo digestion. This reveals a critical non-amyloidogenic function of APP in maintaining tissue homeostasis through efficient efferocytosis, with broad implications for inflammatory and neurodegenerative disorders that warrant further investigation.

## Introduction

Efferocytosis, the recognition, engulfment, and degradation of apoptotic cells (ACs) by phagocytes, is a fundamental biological process essential for tissue homeostasis, development, and resolution of inflammation ^1^. Efficient efferocytosis prevents the accumulation of dying cells, which would otherwise undergo secondary necrosis and release pro-inflammatory intracellular contents, triggering chronic inflammation and autoimmune responses. Failure of efferocytosis has been implicated in a wide range of pathological conditions, including autoimmune disorders, chronic inflammatory diseases, atherosclerosis, and neurodegenerative conditions such as Alzheimer’s disease and Parkinson’s disease ^2^.

After phagocytes engulf ACs, phagosomes fuse sequentially with early endosomes, late endosomes^3^, and ultimately lysosomes, forming acidic phagolysosomes that degrade AC cargo ^4^. Lysosomes are the terminal degradative compartments of the endolysosomal system and are central to efficient clearance of AC. Lysosomal biogenesis, membrane fusion dynamics, and acidification directly determine the speed and efficiency of phagolysosome maturation and AC degradation ^5,6^. Lysosomal biogenesis is transcriptionally regulated by the master regulator transcription factor EB (TFEB) and the coordinated lysosomal expression and regulation (CLEAR) gene network, which controls the expression of lysosomal membrane proteins, hydrolases, and V-ATPase subunit ^7^. However, the upstream signals and molecular mechanisms that activate lysosomal biogenesis specifically during efferocytosis remain incompletely understood. Identifying the regulatory pathways that govern lysosomal function during AC degradation is therefore critical for understanding how cells maintain homeostasis and prevent inflammatory disease.

Amyloid precursor protein (APP) is a highly conserved, ubiquitously expressed type I transmembrane glycoprotein with a large extracellular domain, single transmembrane region, and short intracellular tail ^8^. APP is present in both invertebrates, such as *Caenorhabditis elegans* and *Drosophila melanogaster,* and mammals, highlighting its evolutionary conservation and fundamental biological importance ^9,10^. APP undergoes complex proteolytic processing through two competing pathways. In the non-amyloidogenic pathway, APP is cleaved by α-secretase at the cell surface, releasing the soluble ectodomain sAPPα and preventing Aβ formation. This is followed by γ-secretase cleavage, which releases the APP intracellular domain (AICD) into the cytoplasm. In contrast, the amyloidogenic pathway involves sequential cleavage by β-secretase (BACE1) and γ-secretase, primarily occurring in endosomal compartments, which generates Aβ peptides of varying lengths (Aβ40, Aβ42) along with AICD. The balance between these pathways is critically influenced by APP subcellular localization and trafficking, with endosomal accumulation of APP favoring amyloidogenic processing.

While the amyloidogenic processing of APP and its role in AD pathogenesis have been extensively studied, the physiological functions of APP remain incompletely understood. Investigating APP’s endogenous function is experimentally challenging due to the presence of three closely related APP family members in vertebrates: APP, APLP1, and APLP2 ^9^. Therefore, the analysis of APP function in *Drosophila* offers a promising alternative. The *Drosophila appl* (amyloid protein precursor-like) gene encodes the sole fly homolog of the mammalian APP family, providing a simplified genetic system for dissecting APP function without the confounding effects of functional redundancy ^11^. Recent studies have demonstrated that loss of APPL in *Drosophila* leads to neuronal cell death and enlarged early endosomal compartments, and APPL associates with recycling endosomal to modulate protein aggregation, suggesting a conserved role for APP family proteins in maintaining endolysosomal lysosomal integrity ^12,13^. The secreted form of APPL (sAPPL) interacts with glia, affecting their endosomes and the engulfment receptor-Draper, which is essential for clearing the neuronal debris ^14^. These findings suggest that APPL is crucial for adult brain homeostasis, mediating neuroglial signaling for neuronal survival and clearance of ACs. Furthermore, APPL overexpression induces apoptosis in wing imaginal discs by inhibiting the NEDD8 conjugation pathway, disrupting SCF ligase functions, and reducing neddylated cullins and SCF activity ^15^. This dual role of APPL in promoting cell survival under normal conditions and inducing apoptosis under certain circumstances highlights the complexity of its physiological functions. Emerging evidence suggests that APP plays important roles in neuronal development, synaptic function, axonal transport, and cell adhesion. APP has been shown to interact with a diverse array of intracellular proteins, including G proteins ^16^, Fe65 ^17^, and the PIKfyve complex ^18^, implicating it in signal transduction, transcriptional regulation, and endosomal membrane dynamics. Notably, APP localizes to endosomal compartments and has been shown to regulate PI(3,5)P2 vesicle formation through its interaction with PIKfyve, suggesting a broader role in endosomal trafficking and membrane remodeling. However, whether APP or its homologs function cell-autonomously in phagocytes to regulate AC degradation and the molecular mechanisms underlying such a role remain unexplored.

Spastin is a microtubule-severing AAA ATPase encoded by the *spast* gene ^19^. Mutations in *spastin* cause autosomal dominant spastic paraplegia type 4 (SPG4), the most common form of hereditary spastic paraplegia (HSP), a neurodegenerative disorder characterized by progressive lower limb weakness and spasticity due to defective axonal transport ^20^. Spastin severs microtubules by recognizing and binding to the microtubule lattice through its AAA ATPase domain, generating mechanical force that disrupts tubulin–tubulin interactions ^21^. Spastin contains an N-terminal microtubule-interacting and trafficking (MIT) domain that mediates interactions with the endosomal sorting complex required for transport-III (ESCRT-III) proteins, including IST1 and CHMP1B ^22^. Through its interaction with ESCRT-III, Spastin is recruited to endosomal membranes, where it severs microtubules to facilitate endosomal tubule fission, a critical step in the biogenesis of multivesicular bodies (MVBs) and lysosomes. Loss of Spastin function impairs endosomal tubule fission, leading to the accumulation of elongated endosomal tubules, defective lysosomal biogenesis, and impaired degradative capacity ^22^. In the context of neurodegeneration, spastin dysfunction has been linked to endosomal abnormalities and lysosomal defects. Interestingly, spastin-mediated microtubule severing has also been implicated in Alzheimer’s disease pathology, where tau-induced mislocalization of tubulin-modifying enzymes leads to excessive microtubule severing and dendritic spine loss ^23^. However, the potential connections between Spastin function, endolysosomal homeostasis, and the clearance of cellular cargo, including apoptotic cells, remain largely unexplored.

While APP has been extensively studied in the context of Alzheimer’s disease and amyloid production, its non-amyloidogenic physiological functions, particularly in cellular degradative pathways, have received limited attention. Similarly, although Spastin’s role in endosomal dynamics is recognized, its potential involvement in the degradation of phagocytic cargo has not been investigated. A critical gap exists in our understanding of how APP might regulate endolysosomal homeostasis and whether this function is relevant to efferocytosis. Given that both proteins localize to endosomal compartments and that endolysosomal dysfunction is a common feature of failed efferocytosis and neurodegenerative disease, investigating a potential APP-Spastin axis in regulating apoptotic cell degradation represents a significant opportunity to uncover novel physiological functions of APP and to better understand the cellular mechanisms underlying tissue homeostasis.

In this study, we performed quantitative proteomics on *Drosophila* macrophage-like S2 cell supernatants to identify secreted bridge molecules linking dying cells to phagocytes. Unexpectedly, we also detected APPL, a membrane-associated protein, and later found that its intracellular domain—not its secreted form—is essential for AC degradation in *Drosophila*, indicating its non-amyloidogenic role. Using immunoprecipitation-mass spectrometry (IP-LC MS/MS), we identified a direct interaction between APPL and Spastin mediated by the Spastin MIT domain. This interaction promotes microtubule fragmentation and is essential for endosomal tubule fission and the conversion of late endosomes to lysosomes. Loss of APPL impairs Spastin-dependent microtubule severing, disrupts lysosomal enzyme trafficking, and results in fewer dysfunctional lysosomes with reduced acidity and proteolytic activity, thereby impairing the degradation of ACs. We demonstrated the evolutionary conservation of the APPL-Spastin axis in *C. elegans* and mice. Our findings establish an evolutionarily conserved APP-Spastin pathway governing lysosomal biogenesis and apoptotic cargo digestion, revealing a critical non-amyloidogenic role for APP in maintaining tissue homeostasis through efficient efferocytosis. These results have important implications for understanding the APP’s physiological functions of APP and the pathogenic mechanisms underlying inflammatory and neurodegenerative disorders, including AD, associated with defective efferocytosis.

## Results

### APPL regulates efferocytosis both in S2 cells and *Drosophila* macrophages

To identify secreted proteins involved in efferocytosis, we first established a dose-dependent efferocytosis assay in S2 cells. Croquemort (Crq) is a well-described engulfment receptor, and its expression can be regulated by changes in apoptosis levels in embryonic macrophages ^24^. During the process of efferocytosis in S2 cells, *crq* expression in a macrophage-like lineage derived from late embryonic stages (2) was time- and dose-dependent. Consistent with previous studies, we found a dose-dependent increase in *crq* expression when macrophages engulfed increased amounts of ACs for 6 h **(Fig. 1A)**. Based on transcriptome analysis, we identified some regulatory factors responsible for ACs clearance, such as Wun2 ^25^. During efferocytosis, "Eat-Me" signals are exposed and recognized by phagocytes either directly through phagocyte receptors or indirectly through secreted proteins that function as bridge molecules that cross-link dying cells to phagocytes, such as TTR-52 ^26^ and Gas6 ^27^. However, screening for secretory proteins that regulate efferocytosis remains a major challenge. Thus, to identify the secreted proteins potentially involved in efferocytosis, we performed quantitative proteomic analysis of the supernatant of S2 cells that had engulfed a single or double dose of ACs for 6 h, as well as S2 cells that were not exposed to ACs **(Fig. 1B)**. A total of 127 proteins were upregulated in a dose-dependent manner **(Supplementary Table 1)**. Notably, this screening method identified three secreted proteins—Pretaporter, DmCaBP1, and Calreticulin—that have previously been reported to be involved in the clearance of ACs in *Drosophila* ^28–30^, validating the approach. Within this set, our screening approach specifically identified APPL, the sole *Drosophila* homologue of APP **(Fig. 1C)**. We prioritized proteins with FPKM values >1.5-fold upregulation, predicted signal peptides, and no prior functional annotation in efferocytosis, yielding 15 candidates for RNAi (RNA interference) screening **(Supplementary Table 1)**. We individually silenced 15 secretory proteins to confirm the proteomics results and found that eight of the 15 secretory proteins (CG3505, CG11598, CG33126, CG4920, CG5392, CG16712, CG8050, and CG31619) resulted in a significant reduction in efferocytosis efficiency in S2 cells **(Fig. 1D; Supplementary Table 2)**. Meanwhile, *appl* RNA interference (RNAi) cells exhibited a significantly reduced efferocytosis rate **(Fig. 1E)**. APP is a single-pass type I transmembrane glycoprotein with a large extracellular domain, and its specific physiological functions remain unknown. As *Drosophila* APPL was highly expressed in the supernatant of phagocytic S2 cells, this suggests that APPL may play a role in the clearance of ACs. First, we generated *appl^ko^* mutants in S2 cells and flies using CRISPR/Cas9, which led to the early termination of APPL translation **(Fig. 1F)**; both qRT-PCR and immunofluorescence staining confirmed *appl^ko^* as a null allele mutant **(Fig. S1A and S1B)**. Previous research has found that loss of function of APPL can lead to abnormal enlargement of early endosomes, resulting in the death of neurons in the brains of adult fruit flies during the early stage ^12^. Therefore, we counted the number of ACs and macrophages from stage 11 to stage 15 of fly embryos ^31^, and found that ACs were slightly accumulated in *appl^ko^* flies while macrophages showed no significant difference compared to WT **(Fig. S1C and S1D)**, indicating that the absence of APPL may result in a deficiency in AC degradation without affecting macrophage development. As APPL is also essential for neuronal morphology and function in *Drosophila*, to eliminate the excessive ACs produced by other types of cells, we introduced an *appl* RNAi fly that specifically knocked down *appl* in macrophages. We found that AC cargos mainly accumulated inside macrophages at stage 15 of embryo development **(Fig. 1G to 1I)**. Our results demonstrated that APPL regulates efferocytosis both *in vitro* and *in vivo*.

**Fig. 1.**
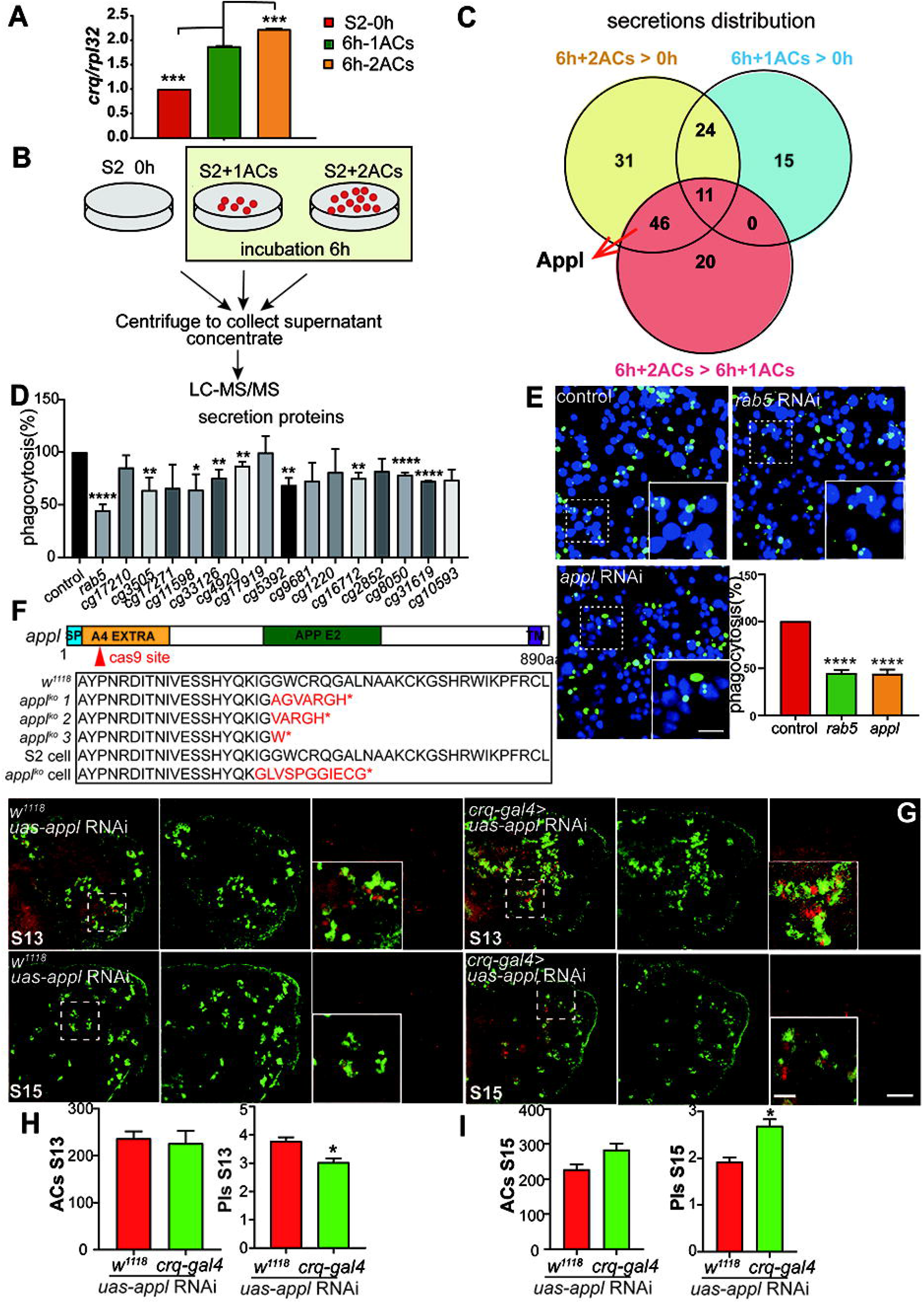
APPL regulates efferocytosis in *Drosophila melanogaster*. **(A)**. The *crq* mRNA level was induced in a dose-dependent ACs amount manner in S2 cells. **(B)**. Schematic representation of the samples used for LS-MS/MS analysis. We collected the supernatant from S2 cells that ingested different doses of ACs, and three samples: S2 cells without ACs, S2 cells with one dose of ACs, and S2 cells with two doses of ACs were used for the proteomic analysis. **(C).** Venn diagram showing the differentially expressed secretory proteins among the three groups. APPL was included in 6h+2ACs>S2 and 6h+2ACs>6h+1ACs. **(D)**. Fifteen secretion proteins were selected for RNAi treatment in S2 cells, and phagocytosis was analyzed in three independent experiments; Data are shown as mean ± SEM. \**p* < 0.05; \*\**p* < 0.01; \*\*\*\**p* < 0.0001. (one-way ANOVA, Dunnett’s multiple comparison test). **(E)**. Confocal images showing ACs efferocytosis by control, r*ab5* RNAi-, and *appl* RNAi-treated S2 cells. Live cells are shown in blue, and FITC-labeled ACs are shown in green. Scale bars: main fields, 50 μm; insets, 10 μm. The column diagram indicates the statistical results of efferocytosis in **(E)**. three independent experiments were repeated, data are shown as mean ± SEM; \*\*\*\**p* < 0.0001. (one-way ANOVA, Dunnett’s multiple comparison test). **(F).** Schematic representation of the CRISPR/cas9 strategy used for the *appl* genome. The red triangles indicate the *appl* sgRNA target sites. Sequencing results showed that the fly *appl^ko^* mutants expressed an early terminated APPL protein, and *appl^ko^* cells were terminated at 113aa. **(G)**. Stage 13 and 15 embryos of *w^11^*^18^/*uas-appl* RNAi and *crqgal4*> *UAS-appl* RNAi flies were stained with anti-Crq (green) and 7-AAD (red). Scale bars, main fields: 100 μm; insets: 20 µm. **(H)**. The column diagram indicates the statistical results for the number of ACs per embryo and PIs at stage 13 in *w^11^*^18^/*uas-appl* RNAi and *crqgal4> uas-appl* RNAi flies. At least three independent experiments were performed, and five embryos were counted in each experiment. Data are shown as mean ± SEM, \**p* < 0.05 (Student’s two-tailed unpaired t-test). **(I)**. The column diagram indicates the statistical results for the number of ACs per embryo and PIs at stage 15 in *w^11^*^18^/*uas-appl* RNAi and *crqgal4> UAS-appl* RNAi flies. At least three independent experiments were performed, and five embryos were counted for each experiment. Data are shown as mean ± SEM, \**p* < 0.05 (Student’s two-tailed unpaired t-test).

### The internalization domain of APPL is essential for AC degradation in S2 cells

APP has mainly been investigated in connection with its role in AD because of its cleavage, resulting in the production of Aβ peptide that accumulates in plaques characteristic of this disease ^32^. APP is an evolutionarily conserved protein found in humans and many other species, including *Drosophila*, suggesting its important physiological functions. In addition to Aβ, several other fragments are produced by the cleavage of APP, including large secreted fragments derived from the N-terminus and small intracellular C-terminal fragments ^9^. In contrast to mammals, which express three APP family members, *Drosophila* expresses only one APP protein, APPL. As the subcellular localization of APP varies in different cell types, APPL is mainly expressed in cytoplasmic vesicles in S2 cells **(Fig. 2A, Fig. S2)**. Similar to APP, APPL is processed by several secretases, resulting in the secretion of fragments and the generation of intracellular AICD. We confirmed that mature APPL was secreted by S2 cells using a GFP nanobody capture assay ^33^**(Fig. 2B)**.

**Fig. 2.**
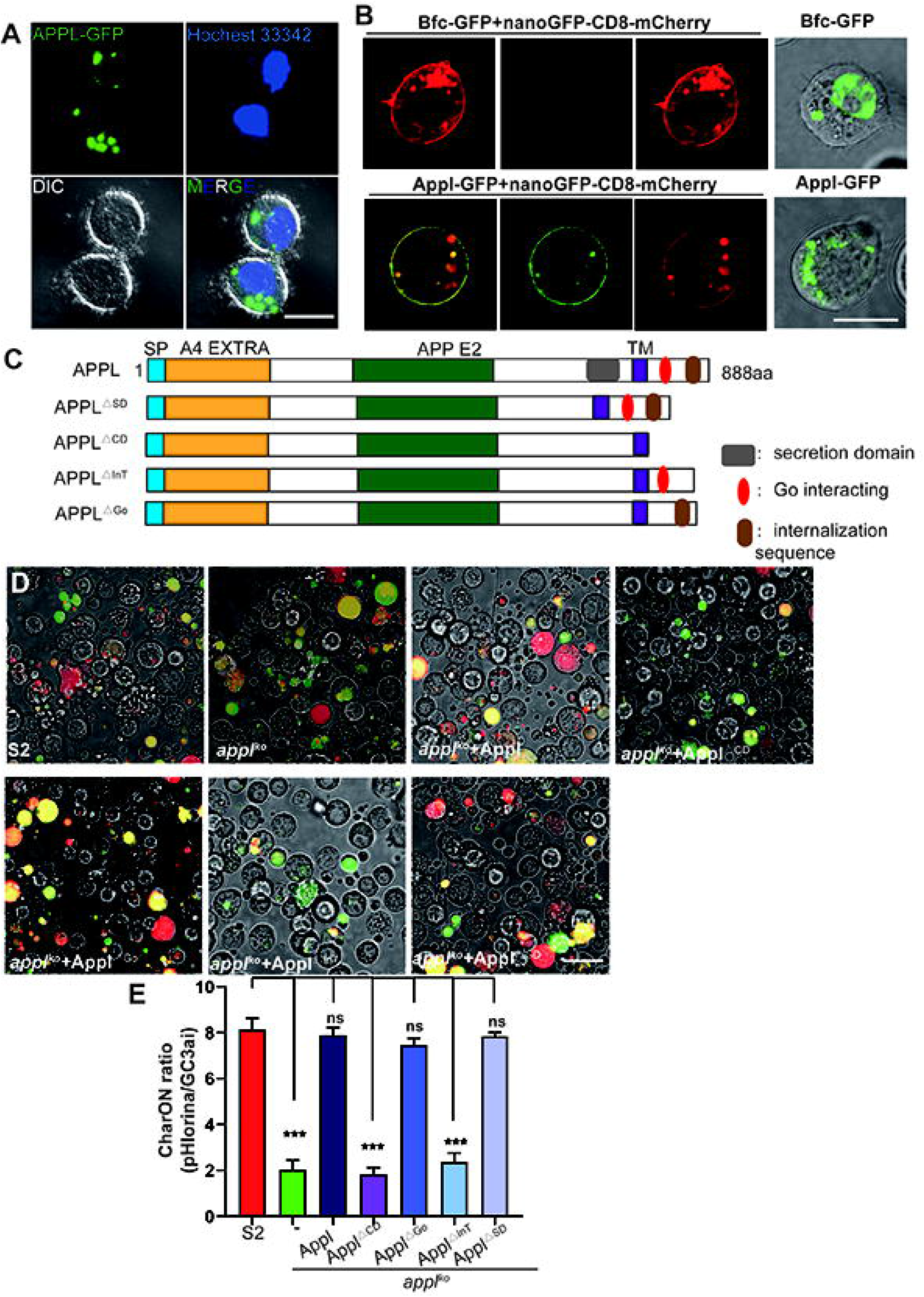
APPL is a secreted protein, and the internalization sequence is essential for efferocytosis of APPL. **(A)**. APPL was localized in the cytoplasm of S2 cells. S2 cells expressing APPL with C-terminal GFP were stained with Hochest 33342 before observation using confocal microscopy, scale bars, 10 μm. **(B)**. APPL is secreted by S2 cells. Transfect APPL-GFP and nanoGFP-CD8-mCherry into S2 cells, respectively, after 48h transfection, transfer the supernatant of S2 cells transfected with APPL-GFP into S2 cells transfected with nanoGFP-CD8-mCherry (removing the supernatant) for 12 h, GFP fluorescence images were observed in confocal microscopy. Scale bars, 10 μm. The nuclear-localized protein Bfc-GFP was used as a negative control. **(C)**. Schematic representation of APPL truncation. APPL^△SD^ represents APPL lacking secretion domain; APPL^△CD^ represents APPL lacking the entire cytoplasm domain; APPL^△InT^ represents APPL lacking the internalization sequence; APPL^△Go^ represents APPL lacking the Go-interacting domain. **(D)**. Confocal images showing APPL rescue of AC degradation in *appl^ko^* cells. *appl^ko^* cells were transfected with different APPL truncations for 48 h and observed by confocal microscopy together with S2 and *appl^ko^* cells. The CharON stable S2 cell line, which expressed a caspase-activated GFP and a pH-insensitive mApple (pHlorina), was treated with 0.25 μg/mL actinomycin D for 6 h, and then added to S2, applko and *appl^ko^* cells expressing different APPL truncations, respectively, and observed by confocal fluorescence microscopy. Scale bars, 20 μm. **(E)**. The column diagram shows the ratiometric CharON (pHlorina/GC3ai) signal during macrophage engulfment. At least three independent experiments were performed, data are shown as mean ± SEM, **\*\*\****p* < 0.001 (one-way ANOVA, Dunnett’s multiple comparison test).

Previous research has found that the secreted form of APPL is important for fly neurite length ^11^; therefore, we expressed a secretion-deficient form (APPL^△sd^) ^34^ in *appl^ko^* cells to verify whether the secretion of APPL was required for efferocytosis as well **(Fig. 2C)**. We then induced a cell line named CharON, derived from a previous report that specifically labeled positive ACs and engulfed ACs (by expressing a vector carrying GC3AI and pHlorina: ACs first turn GFP, and the intensity of pHlorina increases as the GFP gradually quenches after engulfment and acidification) ^35^, to mimic the process by which ACs are engulfed and degraded by macrophages. We added apoptotic CharOn cells to fresh S2 and *appl^ko^* cells, and the *appl^ko^* cells showed a clear defect in degradation **(Fig. 2D).** Although APPL^△sd^ was not successfully secreted from S2 cells, it could still rescue the phagocytic defect phenotype of *appl^ko^* cells **(Fig. 2D and 2E)**, indicating that APPL’s role in efferocytosis is independent of its secreted form. Correspondingly, the secreted form of APPL failed to rescue the degradation function in *appl^ko^* cells (**Fig. S3A to S3C)**, whereas APPL rescued the deficiency of AC degradation in S2 cells (**Fig. S3D)**, confirming that the intracellular domain is both necessary and sufficient. The cytoplasmic domain of APPL contains a Goα-binding site and an internalization domain. The Goα-binding site mediates downstream signal transduction ^36^, and the internalization domain contains a highly conserved YENPTY motif that regulates vesicle transport and endocytic sorting of membrane proteins ^37^. We expressed APPL^△Go^ and APPL^△InT^ in *appl^ko^* cells, respectively, and found that the internalization domain of APPL was essential for the degradation of ACs **(Fig. 2D and 2E)**, indicating that APPL regulates ACs degradation by mediating protein transport and localization.

### APPL interacts with Spastin to regulate the length and stability of microtubules

To further clarify the regulatory mechanism of APPL, we performed IP-LC MS/MS to screen for proteins that interact with APPL. We expressed APPL-HA in S2 cells and performed immunoprecipitation followed by mass spectrometry in *Drosophila* S2 cells **(Fig. 3A and 3B)**. Among the 149 candidates **(Supplementary Table 3)**, we screened 10 intermembrane proteins and identified CG5977-Spastin (Spas) **(Supplementary Table 4)**, a member of the AAA ATPase family that assembles into hexamers and severs microtubules along their length **(Fig. 3C)** ^38^. Microtubules are dynamic cytoskeletal polymers that play central roles in cell division, intracellular transport, cell migration, signaling, and morphogenesis ^39^. The morphology and dynamics of microtubule networks are important for their versatile cellular functions. To validate this interaction, we performed proximity labeling (TurboID) in S2 cells and detected a reciprocal interaction between Spas-Flag and HA-APPL **(Fig. 3D)**, and the BiFC assay of Spas and APPL in 293T cells confirmed this interaction **(Fig. S4A)**, indicating that APPL and Spas function together to regulate efferocytosis. Correspondingly, *spas* RNAi treatment in S2 cells showed a reduced degradation ratio **(Fig. 3E and 3F)**. There are two key functional domains in Spas: the MIT domain, which represents Spas recruitment to endosomal membranes and relies on the interaction with other proteins, and the AAA domains, which associate to form a hexameric ring structure with catalytic activity ^40^. To identify the Spas region responsible for APPL binding, we generated truncated constructs corresponding to the MIT (Spas-1) and AAA (Spas-2) domains **(Fig. 3G).** TurboID assays ^41^ revealed that only the MIT domain directly binds to APPL **(Fig. 3H)**, demonstrating that the MIT domain of Spas may play an essential role in macrophage efferocytosis. As the internalization domain of APPL contains a highly conserved YENPTY motif that regulates vesicle transport and endocytic sorting of membrane proteins, we performed APPL^ICD^ and Spas, and verified that the APPL internalization domain was essential for this interaction **(Fig. S4B and S4C)**. Spas promotes the division of endosomal tubes, and its deficiency leads to incorrect transport of lysosomal enzymes and abnormalities in downstream lysosomes. As APPL interacts with the MIT domain of Spas and is required for cargo degradation, we speculated that APPL is associated with microtubule cutting. When we co-expressed Spas-mCherry and β-tubulin-mStaysGold in S2 and *appl^ko^* cells, respectively, Spas overexpression resulted in excessive fragmentation of microtubules in S2 cells, whereas long microtubules were still observed in *appl^ko^* cells **(Fig. 3I)**. Except for Spas, there are two other MT-severing enzymes: Katanin and Fidgetin, Co-IP assay proved APPL had no interaction with Katanin and Fidgetin, and the *appl* deletion do not affect their cutting of microtubules **(Fig. S4C to S4G)**, further confirming the specific regulatory mechanism of APPL-Spas. Next, microtubule morphology was similar in S2 and *appl^ko^* cells when Spas lacking the MIT domain were expressed **(Fig. 3I and 3J)**. These results indicate that the interaction between APPL and Spas is important for microtubule cutting, which ultimately regulates AC degradation in macrophages, though the precise mechanistic link requires further investigation.

**Fig. 3.**
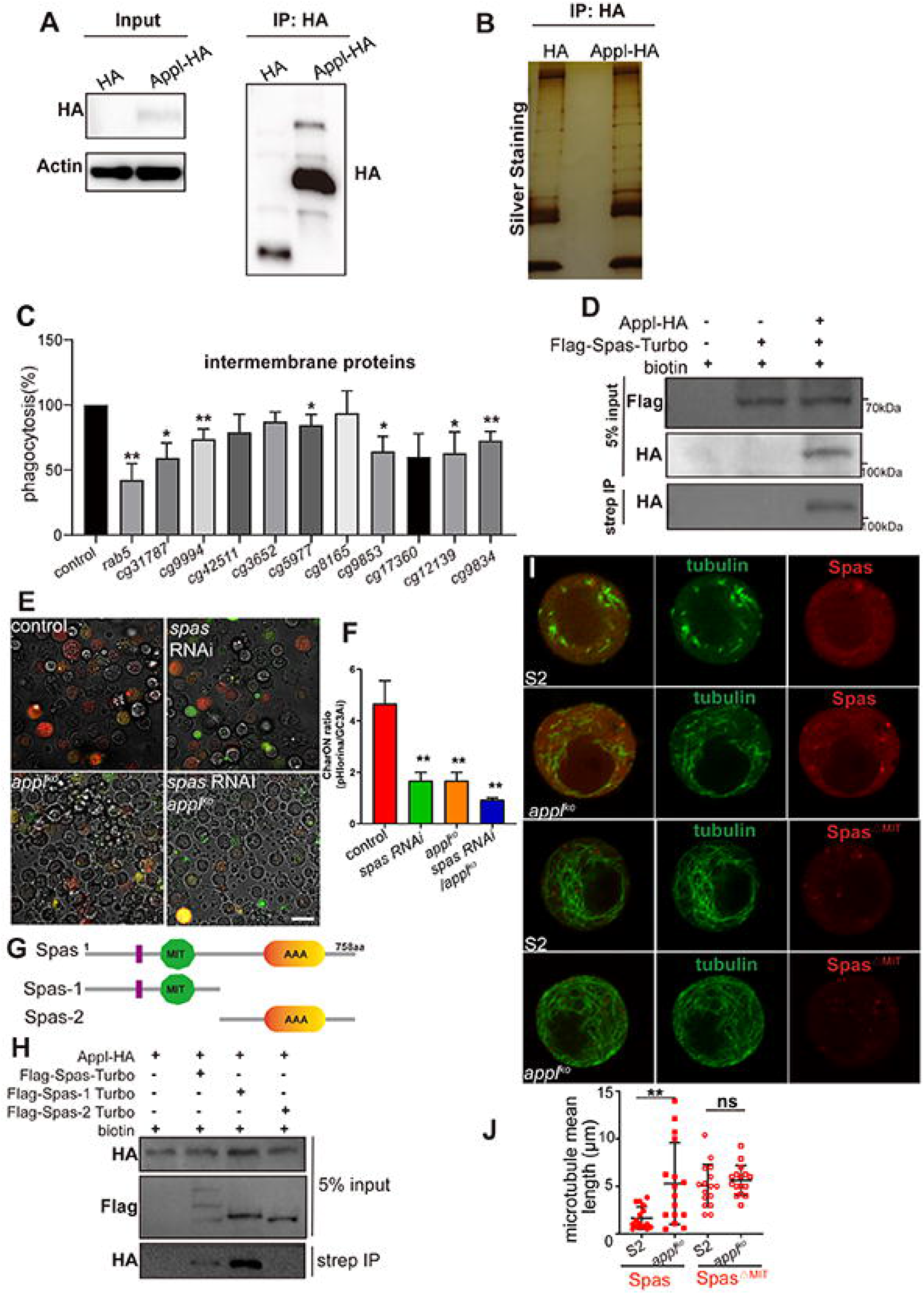
APPL interacts with Spastin to regulate microtubule length. Immunoblot analysis **(A).** and silver staining **(B) of the same samples.** showed the expression of APPL-HA in S2 cells, and APPL and proteins interacting with APPL were enriched using anti-HA magnetic beads. **(C).** Ten intermembrane proteins were selected for RNAi treatment in S2 cells, and efferocytosis was analyzed in three independent experiments; data are shown as mean ± SEM, \**p* < 0.05; \*\**p* < 0.01(one-way ANOVA, Dunnett’s multiple comparison test). **(D).** The TurboID assay revealed that Spastin interacts with APPL. HA-APPL and Flag-Spastin-TurboID were co-transfected into S2 cells, and the cells were treated with 50 μM biotin for 10 min before harvest. Cell lysates were immunoprecipitated using streptavidin magnetic beads and analyzed using Western blotting with anti-HA. **(E)**. Confocal images showing AC degradation by control, *spastin* RNAi, *appl^ko^* and *appl^ko^/spastin* RNAi cells. The CharON stable S2 cell line, which expressed a caspase-activated GFP and a pH-insensitive mApple (pHlorina), was treated with 0.25 μg/mL actinomycin D for 6 h, and then added to S2, *spastin* RNAi-, *appl^ko^* and *appl^ko^/spastin* RNAi cells, respectively, and observed by confocal fluorescence microscopy. Scale bars, 20 μm. **(F)**. The column diagram indicates the statistical results of efferocytosis in **(E)**; at least three independent experiments were performed. Data are shown as mean ± SEM, ** *p*<0.01 (one-way ANOVA, Dunnett’s multiple comparison test). **(G)**. Schematic representation of spastin truncation. Spastin has an N-terminal MIT domain and a C-terminal microtubule-severing AAA ATPase domain. **(H)**. The MIT domain of spastin interacts with APPL. HA-APPL and Spastin truncation -Flag-TurboID were co-transfected into S2 cells, and the cells were treated with 50 μM biotin for 10 min before harvesting. Cell lysates were immunoprecipitated using streptavidin magnetic beads and analyzed using western blotting with an anti-HA antibody. **(I)**. Tubulin with C-terminal mstaygold and Spastin with C-terminal mCherry were co-expressed in S2 and *appl^ko^* cells, respectively. Tubulin with C-terminal mstaygold and Spastin (lacking MIT domain) with C-terminal mCherry were co-expressed in S2 and *appl^ko^* cells, respectively. Scale bars, 5 μm. **(J)**. The mean length of the microtubules per cell in **(I)** was measured using the Fiji software (Analyze Skeleton). Sixteen cells were counted. Data are shown as mean ± SEM, \**p* < 0.05 (Student’s two-tailed unpaired t-test).

### The deficiency of *appl* resulted in insufficient conversion of late endosomes to lysosomes

Spas is recruited to endosomal membranes through the interaction of its MIT domain with CHMP1B of the ESCRT-III complex ^42^. Once recruited, Spas severs microtubules attached to recycling tubules, allowing the tubules to separate from the endosomal membrane and form independent vesicles. BiFC and Y2H assays confirmed the direct interaction between *Drosophila* Spas and Chmp1 **(Fig. 4A and 4B)**, whereas Spas-Chmp1 co-localization (a spatial readout) was unchanged in *appl^ko^* cells **(Fig. 4C and 4D)**. When we detected the co-localization of Spas and various endosomes (Rab5 for early endosomes, Rab7 for late endosomes, and LAMP1 for lysosomes) in S2 and *appl^ko^* cells, respectively, we found that Spas was more localized on the late endosomes in *appl^ko^* cells and less localized on the lysosomes in *appl^ko^* cells in contrast **(Fig. S5A to S5F)**. Furthermore, *appl^ko^* cells showed significantly fewer LAMP1-positive lysosomes with less overlapping Spas **(Fig. S5C)**, which seemed to have a similar phenotype to that of the *spas* mutation. As Spas participates in endosomal tubule fission to mediate the formation of multivesicular bodies, our results indicate that Spas may function in the conversion of late endosomes into lysosomes. Therefore, we observed the co-localization of Spas, Chmp1, and Rab7 in S2 and *appl^ko^* cells and found that Spas and Chmp1 were more specifically located on the periphery of late endosomes in applko cells **(Fig. 4E to 4G)**. To further confirm whether the absence of APPL prevents Chmp1 from dissociating Spas on the late endosomal membrane, we performed the inducible-CuSO4 BiFC assay in S2 cells and the Y3H assay, which indicated that the absence of APPL led to an enhanced interaction between Spas and Chmp1 **(Fig S5G to S5J)**. The increased retention of Spas on Rab7-positive late endosomes and the enhanced interaction between Spas and Chmp1 suggest that APPL regulates complex dynamics rather than recruitment.

**Fig. 4.**
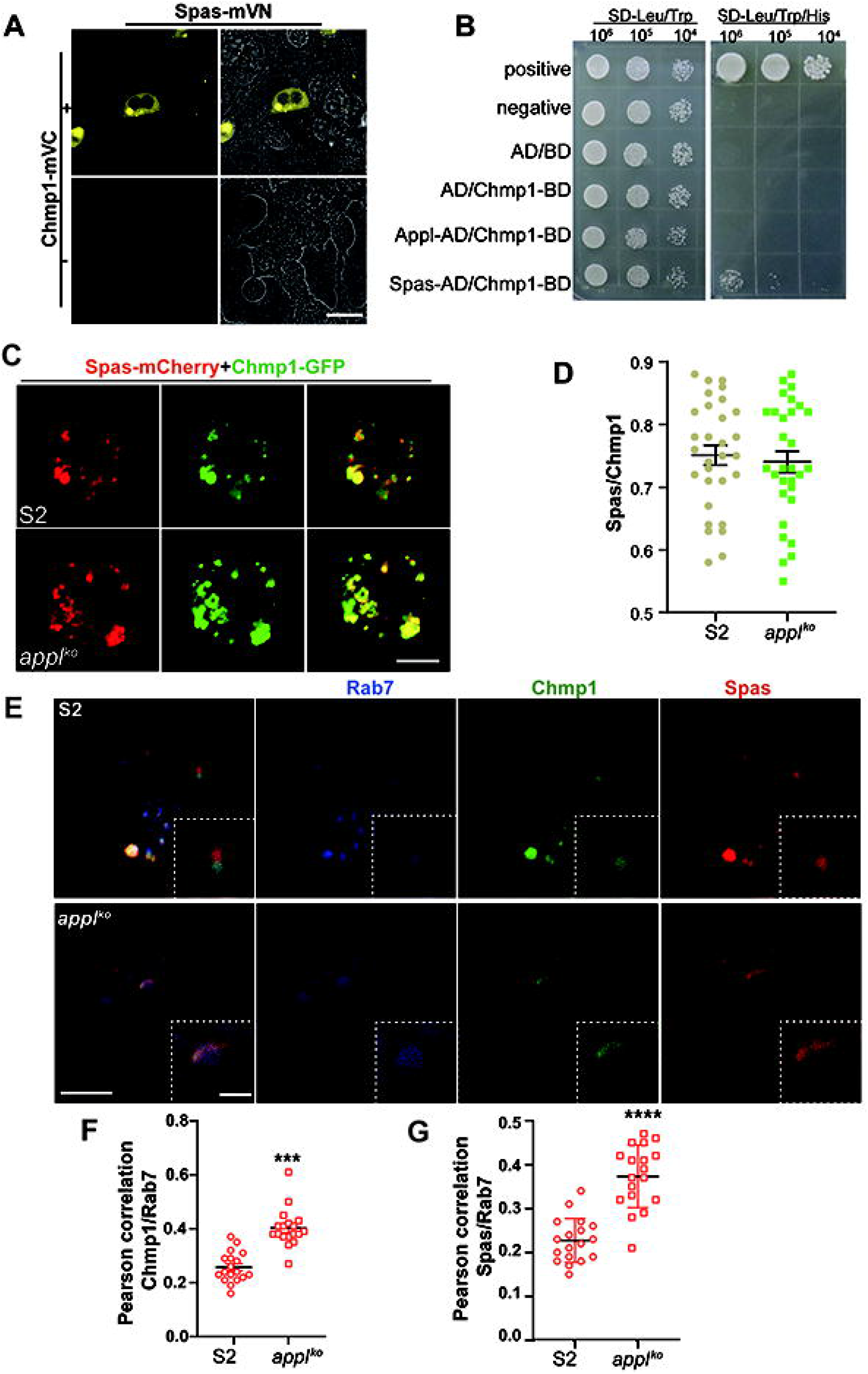
APPL interacts with Spas to regulate the conversion of late endosomes into lysosomes. **(A)**. BiFC showed Spas binding to Chmp1 in 293T cells. Spas-mVN and Chmp1-mVC were co-transfected into 293T cells, and mVenus fluorescence images were captured by confocal microscope. Scale bars, 20 μm. **(B)**. Yeast two-hybrid assays were used to detect interactions between APPL and Chmp1, Spastin, and Chmp1 transformants. Different concentrations of labeled yeast transformants were assayed on SD-His-Trp-Leu plates for growth. Empty AD and Chmp1-BD were used to detect self-activation of the probe. **(C)**. Co-localization of Spas-mCherry and Chmp1-GFP in S2 and *appl^ko^* cells. Scale bars, 5 μm. **(D)**. The extent of co-localization between Spas and Chmp1 proteins was estimated by calculating the Pearson’s correlation coefficient for red and green pixels in each cell using Fiji software (n = 30 cells, *n* = 10 cells in each independent experiment); data are shown as mean ± SEM (Student’s two-tailed unpaired t-test). **(E)**. Co-localization of Spas-mCherry, Chmp1-GFP and Rab5-mTafBFP2 in S2 and *appl^ko^* cells. Scale bars, 5 μm. **(F and G)**. The extent of co-localization between Spastin and Rab7, Chmp1 and Rab7 were estimated by calculating the Pearson’s correlation coefficient for red and blue, green and blue pixels respectively in each cell using Fiji software (n = 30 cells, *n* = 10 cells in each independent experiment); data are shown as mean ± SEM (Student’s two-tailed unpaired t-test).

Defective endosomal tubule fission following SPAS depletion results in the missorting of receptors (including the mannose 6-phosphate receptors (M6PRs)) that traffic via this tubular-vesicular pathway ^21^, and excess detained receptors in the early endosomal compartment results in an endolysosomal degradative compartment. Our results indicated that APPL did not restrain Spas binding to the ESCRT-III complex; however, endosomal tubule fission and downstream trafficking pathways were still unsuccessful, leading to abnormal lysosomal morphology. During endosomal sorting, the ESCRT-III complex containing cargo forms a tubular structure that is disassembled and recycled by Vps4 ^22^, thereby driving membrane detachment. Therefore, we introduced SNX1 (Sorting nexin 1) to indicate tubular endosomes, and Spas deficiency has been reported to reduce tubular SNX1 ^43^. Consistently, we found that SNX1-positive tubular endosomes were more abundant in *appl^ko^* cells than in S2 cells **(Fig. S6A and S6B)**, which regulates the sorting of late endosomes into lysosomes. If this mechanism fails, the recycling tubules cannot break down normally, leading to the accumulation of excessive and elongated tubular structures on endosomes, thereby disrupting the normal recycling of receptors. Furthermore, the reduced degradation of ACs caused by *spas* RNAi in S2 cells could not be rescued by additional APPL **(Fig. S7)**, which suggested that APPL may participate in lysosome regulation via Spas.

### APPL regulated the generation and degradation functions of lysosomes

As APPL deficiency leads to incorrect transport of lysosomal enzymes and abnormalities in downstream lysosomes, we used a human galectin-3 (hGal3)-expressing vector to label the damaged lysosomes ^44^. As expected, *appl^ko^* cells showed more hGal3-mCherry puncta signals **(Fig. 5A and 5B)**, indicating that abnormal lysosomal sorting increases the damage to lysosomes. As *spas* depletion has been reported to cause abnormal lysosomal morphology and fewer lysosomes ^38^, we examined lysosomal function in *appl^ko^* cells. First, we examined the lysosomal structures using transmission electron microscopy (TEM). In *appl^ko^* cells, the number of lysosomes was much lower than that in S2 cells, without contact with the endoplasmic reticulum **(Fig. 5C, 5D and 5F)**. PI(3,5)P_2_ is proposed to be mainly localized to early endosomes, late endosomes, and lysosomes ^45^, and the PX domain of SnxA in *Dictyostelium* showed highly specific binding to PI(3,5)P_2_ ^46^. The co-localization of 2*PX and VAC14 with APPL showed that APPL likely functions in late endosomes or lysosomes **(Fig. S8A to S8C)**. Deletion of *appl* generated fewer late endosomes than early endosomes **(Fig. S8D to S8G)**. Furthermore, BiFC and Co-IP assays showed a direct interaction between APPL and Rab7, indicating that APPL may be recruited to late endosomes to regulate Spas and Chmp1 **(Fig. S8H to S8J)**. The transcriptional level and localization of the lysosomal gene Mitf showed no difference between S2 and *appl^ko^* cells (**Fig. S9A to S9E).** This reduction in the number of lysosomes suggests a defect in their biogenesis or stability, possibly stemming from impaired endosomal tubule fission. We then examined lysosomal acidity and proteolytic activity using Lysotracker Red and Magic Red and found that the intensity of Lysotracker Red and Magic Red was substantially reduced in *appl^ko^* cells **(Fig. 5E and Fig. S9F and S9G)**, suggesting that the loss of APPL affects the acidity and proteolytic activity of lysosomes, which explains why *appl^ko^* cells exhibit defects in AC degradation. Furthermore, we added apoptotic CharOn cells to S2 and *appl^ko^*cells and isolated macrophages from *w^11^*^18^ and *appl^ko^* larvae. Both *appl^ko^* cells *and* macrophages exhibited abnormal AC degradation **(Fig. 5H to 5 K)**, consistent with the lysosomal defects described above, indicating that APPL is required for lysosomal functions. Furthermore, the *appl^ko^* cells presented a much slower degradation speed of *S. aureus* compared to that of S2 cells **(Fig. S9J and S9I)**, indicating that APPL’s role in degradation extends to bacterial cargo in addition to apoptotic cells.

**Fig. 5.**
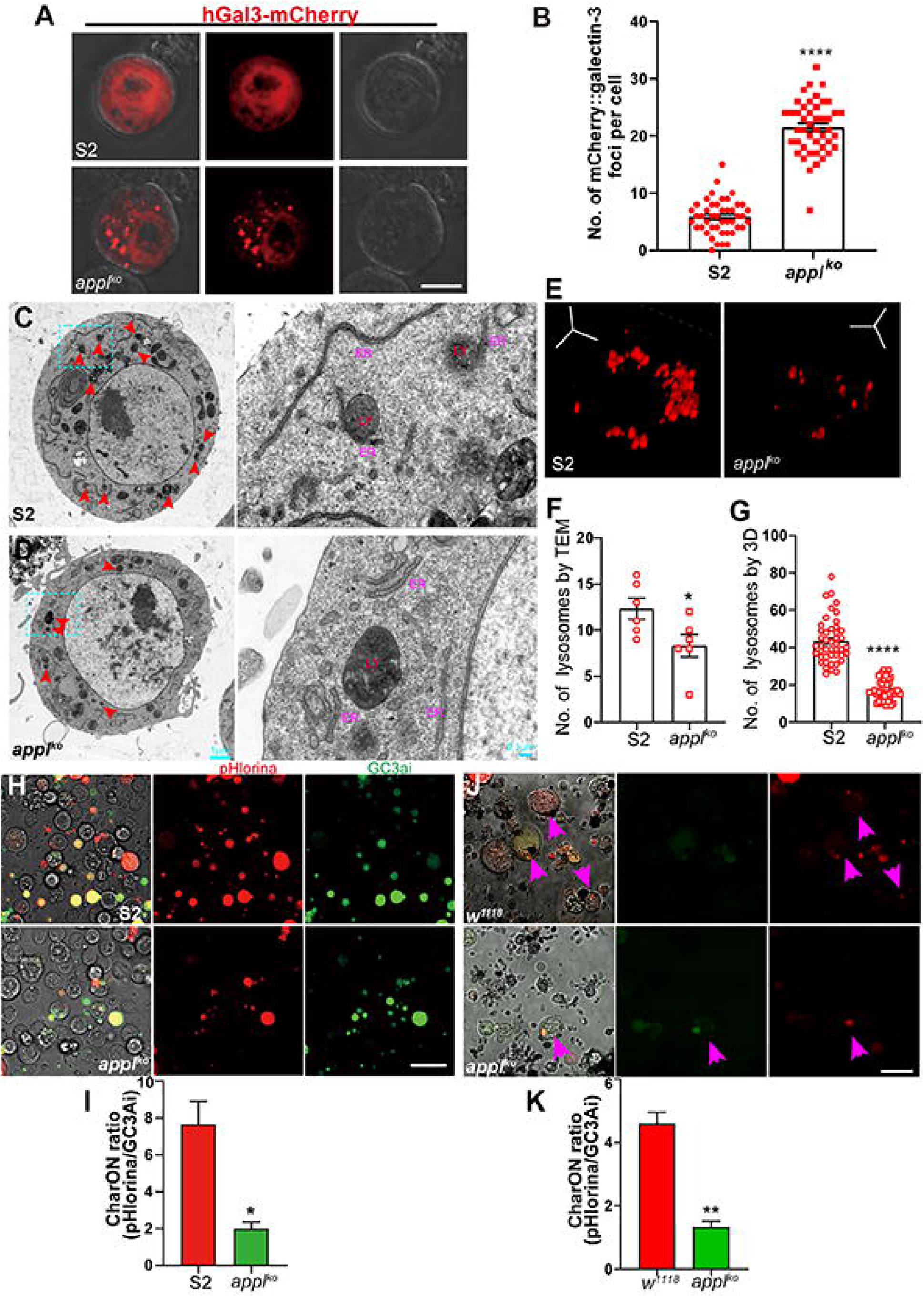
APPL regulates lysosome biogenesis and maturation in macrophages. (A). *appl^ko^* induced lysosomal damage in S2 cells. hGal3 (human Galectin-3) was expressed in S2 and *appl^ko^* cells tagged with mCherry, scale bars, 5 μm. **(B).** The number of hGal3-mCherry foci per cell is shown in the column, n = 15 cells in each of the three independent experiments, data are shown as mean ± SEM, \*\*\**p* < 0.001 (Student’s two-tailed unpaired t-test). **(C and D).** TEM images of lysosomes in S2 and *appl^ko^* cells. Red arrowheads indicate lysosomes, and the regions enclosed by blue rectangles are shown at higher magnification in insets. Insets: ER, endoplasmic reticulum; LY, lysosomes. Scale bars, main fields: 1 μm; insets: 100 nm. **(E).** Confocal fluorescence images of S2 and *appl^ko^* cells stained with LysoTracker Red in the z-axis direction. The number of lysosomes per cell, as determined by TEM is shown in **(F)**. n = 6 cells, data are shown as mean ± SEM; \**p* < 0.05 (Student’s two-tailed unpaired t-test). The number of lysosomes per cell as determined by LysoTracker Red is shown in **(G)**. n = 15 cells in each of three independent experiments, data are shown as mean ± SEM, \*\*\*\**p* < 0.0001 (Student’s two-tailed unpaired t-test). **(H).** Degradation assay in *Drosophila* larval macrophages. The CharON stable S2 cell line, which expressed caspase-activated GFP and a pH-insensitive mApple (pHlorina), was treated with 0.25 μg/mL actinomycin D for 6 h, and then added to isolated macrophages from *w^11^*^18^ and *appl^ko^* larvae, respectively, and observed by confocal fluorescence microscopy. **(I).** The column diagram shows the ratiometric CharON (pHlorina/GC3ai) signal during macrophage engulfment. Three independent experiments were performed, data are shown as mean ± SEM, \**p* < 0.05 (Student’s two-tailed unpaired t-test). **(J).** Degradation assay in S2 cells. The CharON stable S2 cell line, which expressed caspase-activated GFP and pH-insensitive mApple, was treated with 0.25 μg/mL actinomycin D for 6 h, and then added to S2 cells and *appl^ko^* cells, respectively, and observed by confocal fluorescence microscopy. **(K).** The column diagram shows the ratiometric CharON (pHlorina/GC3ai) signal during embryonic engulfment. Three independent experiments were performed. Data are shown as mean ± SEM, \*\**p* < 0.01 (Student’s two-tailed unpaired t-test).

### The mouse App performed a conserved function in AC degradation

The APP family is highly conserved during evolution, with only one APP homolog in either *Drosophila* or *C. elegans*. We found that APL-1 regulates AC degradation via a similar pathway **(Fig. S10; Supplementary Table 5)** as in *Drosophila*. However, APL-1 is extensively expressed in *C. elegans* (especially in non-neural tissues), whereas APPL is restricted to the nervous system, reflecting the high specialization of fruit flies as a model for studying neural behaviors during evolution. To rigorously assess the evolutionary conservation and physiological significance of the non-amyloidogenic functions of APP in mammals, we developed a constitutive *App* loss-of-function mouse model(*C57BL/6J-App*em1) characterized by a precisely targeted exon deletion that eliminates all canonical App isoforms. Comprehensive validation confirmed the complete ablation of App expression at both the transcriptional and protein levels, including in bone marrow–derived macrophages (BMDMs), thereby establishing a bona fide null system **(Fig. 6A to 6C)**. Given the AC cargo degradation function of APPL in *Drosophila*, we assessed the function of App in mammalian cell lines. Using dual-color pH-sensitive CharON Jurkat reporters, which allow for real-time quantitative assessment of both engulfment and intra-lysosomal degradation, we dynamically quantified AC degradation kinetics in primary BMDMs. App deficiency caused a severe, time-dependent impairment in degradative capacity: while wild-type macrophages efficiently degraded >80% of internalized ACs by 12 h post-feeding, *App-/-* BMDMs retained >60% undegraded cargo, as evidenced by persistent green+/red+ puncta **(Fig. 6D and 6E)**. Collectively, these data establish App as an evolutionarily conserved, non-redundant regulator of lysosomal homeostasis in mammalian macrophages and reveal a previously unappreciated amyloid-independent axis through which App governs innate immune tolerance by ensuring efficient efferocytosis, a function whose failure is mechanistically linked to chronic inflammation, autoimmunity, and neurodegeneration. Building on our previous discovery of App-dependent lysosomal defects in S2 cells, we investigated whether this conserved pathway operates in mammalian macrophages. TEM combined with quantitative LysoTracker Red staining and automated image-based lysosome segmentation revealed a reduction in the number of lysosomes in *App-/-* BMDMs, without concomitant changes in autophagosome accumulation or gross cellular morphology, indicating a specific deficit in lysosomal biogenesis or stability **(Fig. 6F to 6I)**. Furthermore, immunoblotting analyses demonstrated a significantly reduced expression of the cation-dependent mannose-6-phosphate receptor (M6PR), a key lysosomal sorting receptor, at both the protein and transcript levels in *App-null* cells **(Fig. 6J and 6K)**. As APP is mainly expressed in neurons and serves as an important component of Alzheimer’s disease, we isolated the cerebral cortex of mutant mice and counted the number of lysosomes in the neurons using TEM. The *App-/-* mice showed a significantly reduced number of lysosomes **(Fig. S11)**, indicating that the function of App in regulating lysosome biogenesis may be related to neurodegenerative diseases. To investigate whether App modulates phagocytic engulfment or the intracellular degradative phase of ACs clearance under physiological conditions *in vivo*, we performed *in vivo* efferocytosis assays using TAMRA-labeled ACs **(Fig. S12)**. Relative to WT controls, *App-/-* mice displayed a profound, highly significant reduction in the percentage of TAMRA⁺ splenic macrophages. In peritoneal macrophages, the proportion of TAMRA⁺ cells were also significantly decreased in *App-/-* mice, although the magnitude of this reduction was smaller than that observed in splenic macrophages, possibly reflecting distinct macrophage subtype-specific requirements for APP or compensatory mechanisms in the peritoneal niche. Because the generation of a robust TAMRA signal is strictly dependent on the lysosomal degradation of ingested ACs, these findings demonstrated that App deficiency impairs the intracellular degradation of ACs but does not affect the initial engulfment step of efferocytosis.

**Fig. 6.**
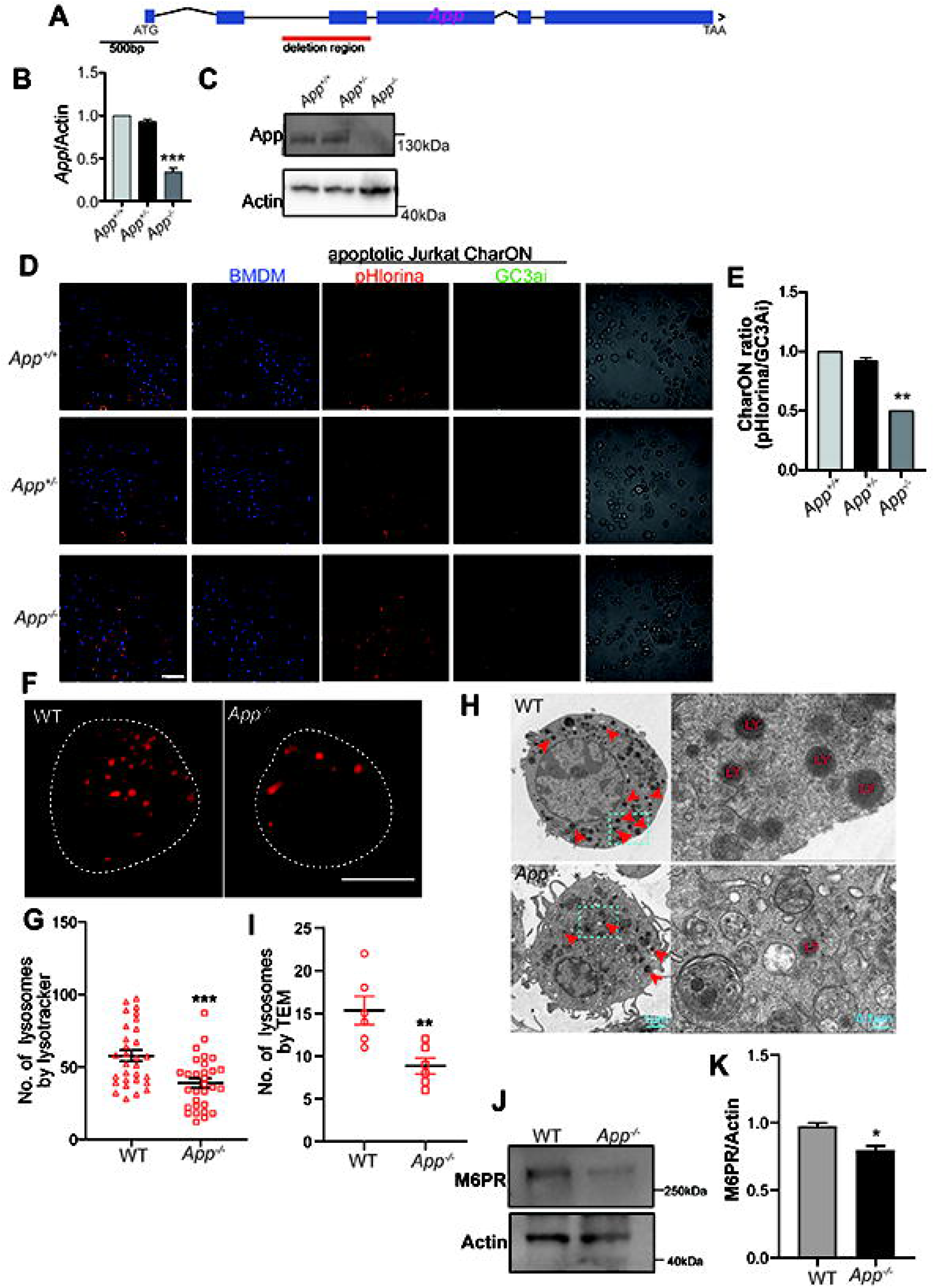
App promotes AC degradation by BMDMs through the regulation of lysosomal quantity and function. **(A)**. Diagram of App protein and targeted deletion strategy. **(B).** *App* mRNA level was analysed by qRT-PCR, *actin* was used as an internal control. BMDMs isolated from *App^+/+^*, *App⁺/⁻* and *App⁻/⁻* were used as samples, and three independent experiments were performed. Data are shown as mean ± SEM. \*\*\**p* < 0.001 (one-way ANOVA, Dunnett’s multiple comparison test). **(C).** Protein was extracted from *App^+/+^*, *App⁺/⁻* and *App⁻/⁻* BMDMs, lysates were analysed using western blot by APP antibody. Actin was used as a loading control. **(D).** Role of APP in the degradation assay of BMDMs. The CharON stable Jurkat cell line, which expressed caspase-activated GFP and pH-insensitive mApple, was treated with 10 ng/mL staurosporine for 12 h, and then added to *App^+/+^*, *App⁺/⁻* and *App⁻/⁻* BMDMs, respectively. BMDMs nuclei were stained with DAPI, and observed by confocal fluorescence microscopy. Scale bars, 50 μm. **(E).** The column diagram shows the ratiometric CharON (pHlorina/GC3ai) signal in **(D).** Three independent experiments were performed. Data are shown as mean ± SEM, \*\**p* < 0.01 (one-way ANOVA, Dunnett’s multiple comparison test). **(F).** Confocal fluorescence images of *App^+/+^* and *App⁻/⁻* BMDMs stained with LysoTracker Red in the z-axis direction. The number of lysosomes per cell as determined by LysoTracker Red is shown in **(G)**. n = 10 cells in each of three independent experiments, data are shown as mean ± SEM, \*\*\**p* < 0.001. (Student’s two-tailed unpaired t-test). **(H)** TEM images of lysosomes in *app^+/+^* and *app⁻/⁻* BMDMs. Red arrowheads indicate lysosomes, and the regions enclosed by blue rectangles are shown at higher magnification in insets. Insets: ER, endoplasmic reticulum; LY, lysosomes. Scale bars, main fields: 1 μm; insets: 100 nm. The number of lysosomes per cell as determined by TEM is shown in **(I)**, n = 6 cells, data are shown as mean ± SEM; ** *p*<0.01. (Student’s two-tailed unpaired t-test). **(J).** M6PR protein levels in WT and *App-/-* cells were analysed using Western blot. Protein was extracted from *App^+/+^* and *App⁻/⁻* BMDMs, and lysates were analysed using western blot by M6PR antibody, Actin was used as a loading control. **(K).** Quantification of the relative protein expression in **(J)**. Three independent experiments were performed, data are shown as mean ± SEM. \*\**p* < 0.01. (Student’s two-tailed unpaired t-test).

### Mammal App regulates lysosome function via interacting with SPASTIN

There are three App members in mammals; however, only App has a significant function in degrading ACs **(Fig. S13A and S13B).** As App shares more conserved functional domains with APPL, we aimed to ascertain whether the secreted extracellular form of App plays a role in efferocytosis within macrophages. Similarly, we engineered a signal peptide-deficient human APP mutant (APP^ΔSP-β^) by precisely excising the canonical N-terminal signal sequence using site-directed mutagenesis. Transfection of HEK293T cells with this construct resulted in the complete abrogation of APP secretion into the conditioned medium **(Fig. 7A)**. Subsequently, to elucidate the compartment-specific requirement for APP in AC degradation, we conducted functional rescue experiments using *App*^–/–^ BMDMs. These cells were transduced with lentiviral vectors expressing either full-length human APP or the non-secreted APP^ΔSP-β^ isoform, which retains only the intracellular β-subunit domain and is thus strictly localized to intracellular compartments, including the endosomes and trans-Golgi network. Notably, both constructs restored the AC degradation capacity to wild-type levels. Quantitative confocal imaging of dual-color CharON Jurkat reporter cells, where green fluorescence indicates acid-sensitive, apoptosis-specific signal degradation within functional lysosomes and red fluorescence marks total internalized cargo, revealed no statistically significant difference in AC degradation efficiency between *App*–/– BMDMs rescued with APP versus APP^ΔSP-β^ **(Fig. 7B and 7C)**. These findings robustly demonstrate that APP contributes to AC degradation independent of its proteolytic shedding or extracellular release. Instead, the intracellular APP pool is necessary and sufficient to support the efficient degradation of ACs. Mechanistically, we investigated whether the intracellular domain of APP regulates lysosomes via SPASTIN in mammalian cells, and Co-IP assays confirmed that SPASTIN interacts with APP **(Fig. 7D; Fig. S13C)**. As SPASTIN regulates membrane remodeling and endosomal fission, we further explored whether SPASTIN contributes to AC degradation in mammalian macrophages and whether this function requires APP expression. To address this, we performed RNAi-mediated knockdown of *spastin* in BMDMs isolated from wild-type (WT) and *App*^⁻/⁻^ mice and quantified AC degradation efficiency over time using the dual-color CharON Jurkat reporter system. Remarkably, *spastin* depletion severely impaired AC degradation in WT BMDMs; however, the same level of knockdown elicited only a marginal defect in *App*^⁻/⁻^ BMDMs **(Fig. 7E and 7F)**, suggesting that most of SPASTIN’s role in AC degradation requires APP, though partial redundancy with other pathways cannot be excluded. These results indicate that SPASTIN and APP function within the same genetic pathway for AC degradation, demonstrating the evolutionary conservation of the APP–SPASTIN regulatory axis during efferocytosis.

**Fig. 7.**
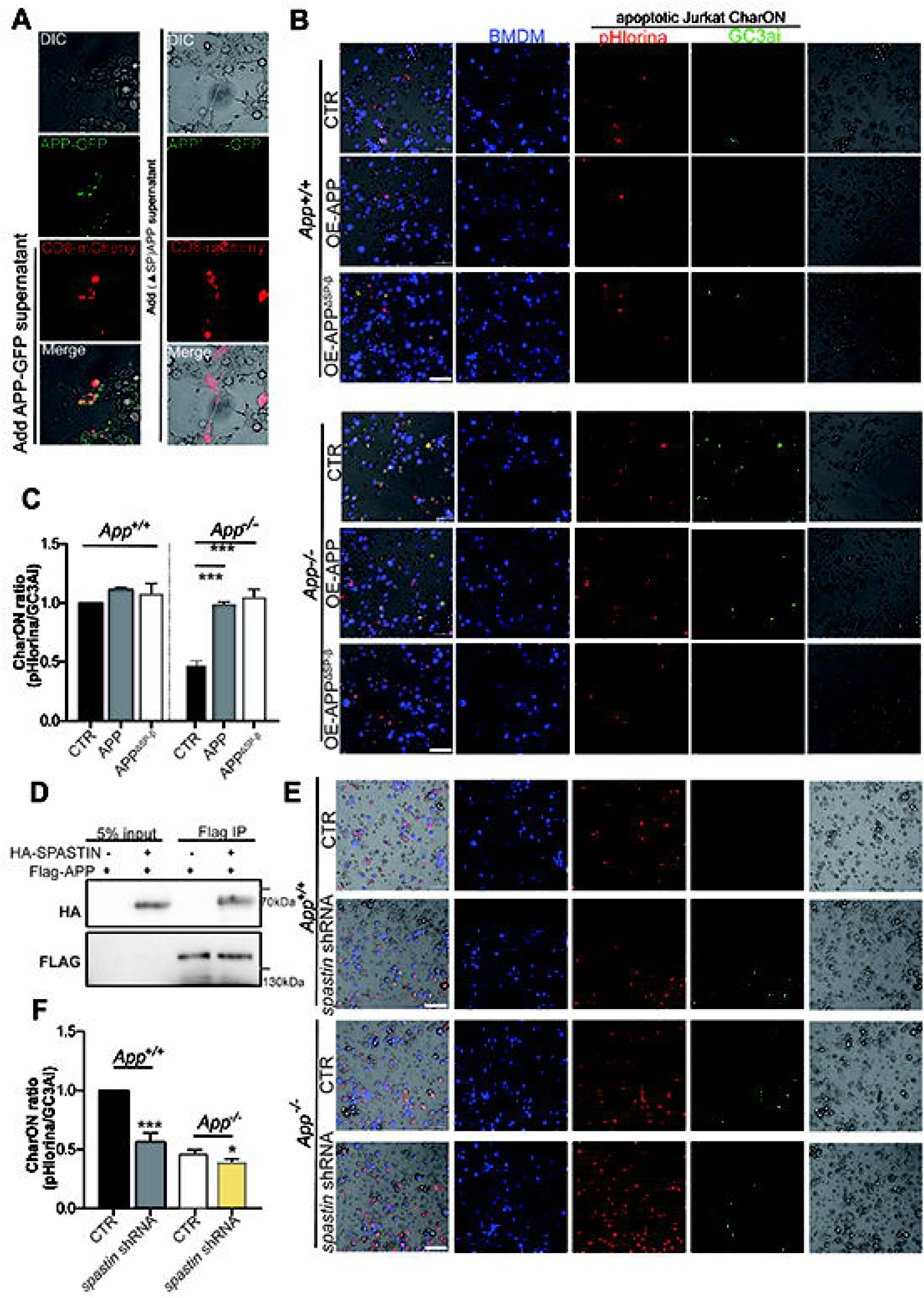
App regulates AC degradation in BMDMs by interacting with SPASTIN via its intercellular domain. **(A)**. APP secretion assay in HEK293T cells. Transfect full-length human APP or a signal peptide-deficient APP construct retaining only the β-subunit (APP^ΔSP-β^) into HEK293T cells, respectively, after 48 h transfection, transfer the supernatant of S2 cells transfected with APP or APP^ΔSP-β^ into HEK293T cells expressing nanoGFP-CD8-mCherry (removing the supernatant) for 12 h, GFP fluorescence images were observed in confocal microscopy. Scale bars, 10 μm. **(B).** The CharON stable Jurkat cell line was treated with 10 ng/mL staurosporine for 12 h, and then added to *App+/+* and *App⁻/⁻* BMDMs re-expressing APP-FL or APP^ΔSP-β^, respectively. BMDMs nuclei were stained with DAPI, and observed by confocal fluorescence microscopy. Scale bars, 50 μm. **(C).** The column diagram shows the ratiometric CharON (pHlorina/GC3ai) signal in **(B).** Three independent experiments were performed. Data are shown as mean ± SEM, \*\*\**p* < 0.001 (one-way ANOVA, Dunnett’s multiple comparison test). **(D).** Co-IP assays in 293T cells to assess the interaction between HA-SPASTIN and FLAG-APP. HA-SPASTIN and FLAG-APP were co-transfected into 293T cells, cell lysates were immunoprecipitated by anti-Flag magnetic beads, and analysed using western blot by anti-Flag and anti-HA. **(E).** *spastin* knockdown impairs AC degradation. The CharON stable Jurkat cell line, which expressed caspase-activated GFP and pH-insensitive mApple, was treated with 10 ng/mL staurosporine for 12 h, and then added to *App+/+* and *App⁻/⁻* BMDMs re-expressing control or *spastin*-targeting shRNA vectors, respectively. BMDMs nuclei were stained with DAPI, and observed by confocal fluorescence microscopy. Scale bars, 50 μm. **(F).** Quantitative analysis of the results presented in **(E)**; Three independent experiments were performed. Data are shown as mean ± SEM, \**p* < 0.05, \*\*\**p* < 0.001 (one-way ANOVA, Dunnett’s multiple comparison test).

## Discussion

While prior work showed APPL affects endosomal morphology and neuronal survival ^12^, our findings establish a previously unrecognized, evolutionarily conserved physiological role for the amyloid precursor protein (APP) and its *Drosophila* homolog, APPL, in regulating endolysosomal homeostasis and degradation of apoptotic cells (ACs) in phagocytes. Using *Drosophila* as our primary model system, we identified a direct interaction between APPL and the microtubule severing ATPase Spastin, mediated by Spastin’s MIT domain. We further demonstrated that APPL regulates the dynamics of the Spastin-Chmp1 interaction on endosomal membranes, as evidenced by enhanced Spastin-Chmp1 binding in APPL-deficient cells, though the precise molecular mechanism by which APPL modulates this interaction requires further investigation. APPL is essential for the dissociation of Spastin from Chmp1 to allow downstream endosome fission, maintaining endosomal tubule fission, lysosome biogenesis, and lysosomal degradative capacity, all of which are indispensable for efficient AC degradation. Critically, we demonstrated that this APPL-Spastin axis is evolutionarily conserved in *C. elegans* (APL-1) and mice (App), where App deficiency similarly impairs AC clearance in macrophages **(Fig. S14)**. These findings reveal a critical non-amyloidogenic function for APP in maintaining tissue homeostasis through efficient efferocytosis, with significant implications for inflammatory and neurodegenerative disorders.

The mechanistic basis for APP’s non-amyloidogenic function lies in its intracellular domain, not its secreted products. Although APPL was detected in the supernatant proteomics, likely due to ectodomain shedding or microvesicle release, our functional rescue experiments in *Drosophila* demonstrated that secretion-deficient APPL(APPL^ΔSD^) fully rescues AC degradation defects in *appl^ko^* cells, whereas the secreted ectodomain alone does not. Similarly, in mammalian cells, a signal peptide-deficient APP construct (APP-^ΔSP-β^) that is not secreted rescues AC degradation as effectively as full-length APP. These findings indicate that APP’s role in efferocytosis depends on its membrane-associated form and internalization domain, which mediates interaction with Spastin. While secreted APPL has been shown to regulate glial endosomes and Draper expression ^14^, our data demonstrate that the intracellular domain is necessary and sufficient for AC degradation in phagocytes, suggesting distinct roles for secreted versus membrane-associated APP in different cellular contexts. The intracellular domain of APP also contains an evolutionarily conserved G-protein-binding motif, enabling APP to function as an atypical G-protein-coupled receptor and perform signal transduction ^36^. AICD directly binds to and activates Gαo (the most abundant G protein in the brain), regulates the PI3K/Akt survival pathway, and mediates the neuroprotective effect of sAPPα in neuronal cells ^16^. However, the Go-binding domain did not affect efferocytosis or degradation. AICD plays a crucial regulatory role in APP endocytic transport, endosome sorting, and signal transduction by interacting with different intracellular proteins. AICD has been proposed to act as a transcriptional regulator through its interaction with the adaptor protein Fe65 and its translocation to the nucleus, forming a complex with histone acetyltransferase (Tip60) ^17^. Hence, identifying proteins that interact with APPL is a key breakthrough point for unveiling the underlying mechanisms.

Using IP-LC MS/MS, we revealed that APPL coordinates microtubule dynamics with endolysosomal membrane trafficking through its direct interaction with Spastin. Previous studies have shown that APP binds to Vac14, a subunit of the PIKfyve complex, thereby enhancing phosphatidylinositol 3,5-bisphosphate [PI(3,5)P₂] synthesis, which is a lipid second messenger essential for endosomal and lysosomal homeostasis and neuronal integrity ^18^. Disruption of the APP-Vac14-PIKfyve pathway leads to aberrant endosomal sorting and neuronal dysfunction. However, our genetic analysis in *Drosophila* demonstrated that APPL does not interact with Vac14 in S2 macrophage-like cells, suggesting the APP-Vac14-PIKfyve pathway may be neuron-specific or mammalian-specific, requiring further investigation.

We identified this alternative pathway as a direct interaction between APPL and Spastin, mediated specifically through the Spastin MIT domain ^42,47^, which operates in phagocytic cells to regulate efferocytosis. This interaction is highly conserved from *C. elegans* to humans, demonstrating that APP and Spastin operate within the same functional pathway for efferocytosis across metazoan evolution. The MIT domain-mediated interaction provides a mechanistic hypothesis for how APP regulates microtubule severing and endosomal membrane remodeling. We propose that APPL-dependent Spastin activity severs microtubules at endosomal tubule necks, facilitating membrane fission. However, direct visualization of microtubule severing and tubule scission in real time will be required to test this model. *spast* mutations are one of the main causes of HSP, and transport disorders in endosomes/lysosomes are often observed in HSP patients ^48^. Membrane scission is an ancient and ubiquitous event during endosomal sorting, where the ESCRT-III subunits CHMP1B and IST1 form a stable scaffold for membrane budding, causing the tube to contract, but they do not cause cutting ^49,50^. Previous reports have found that CHMP1B can bind AAA ATPases-Vps4 to depolymerize and remodel ESCRT-III assembly, leading to membrane contraction and scission ^51^. Spastin is also involved in the scission of recycling endosomal tubules via CHMP1B and IST1^52^, because microtubule disassembly by Spastin and membrane scission mediated by ESCRT-III often occur in combination. We found that APPL does not affect Spastin recruitment to Chmp1-positive endosomes but is required for Spastin-Chmp1 dissociation, as evidenced by enhanced Spastin-Chmp1 interaction in *appl^ko^* cells **(Fig. S5)**. This suggests that APPL modulates the dynamics, rather than the initiation, of the Spastin-ESCRT-III interaction. Some researchers have found that Spastin is recruited to the endosome by CHMP1B, but it does not participate in membrane scission mediated by CHMP1B ^48^. We also found that Spastin colocalized with Chmp1-enriched sites in S2 cells, which increased on late endosomes after *appl* deletion, further confirming that APPL functions in dissociating the complex of Spastin and Chmp1 for endosome fission.

To complete "reverse transport" (recycling sorting receptors back to the Golgi) or form lysosomes, endosomes extend highly dynamic tubular structures (SNX1). These tubes must be cut precisely to release the transport carriers. APP deficiency leads to the abnormal accumulation of tubular SNX1, which is similar to the previously reported phenotypes of *spastin* deletion mutants, indicating that the absence of APP does not prevent the formation of endosomal tubules but blocks their disintegration and recycling. Endosomes continuously extend tubules, which accumulate in APPL-deficient cells. We infer that APPL-dependent Spastin activity normally severs these tubules, as Spastin overexpression fails to fragment microtubules in *appl^ko^* cells However, direct visualization of tubule scission events is needed to confirm this model. Lysosome biogenesis is highly dependent on tubular structures, such as autophagic lysosome regeneration (ALR) pathways, which bud from endosomes and autophagosomes. If the disintegration/recovery of tubular endosomes mediated by APP-Spastin is blocked, the sustained presence of uncleaved tubular endosomes may either physically impede the budding of nascent lysosomes or lead to their aberrant maturation/degradation, resulting in lysosomal degradation dysfunction (such as blocked AC degradation). In addition to Spastin, there are two other MT-severing enzymes: Katanin and Fidgetin ^53^, which have been shown to have no interaction with APPL, and their deletion does not affect their cutting of microtubules, further confirming the specific regulatory mechanism of the APPL-Spastin complex.

Mechanistically, APPL/APP deficiency leads to fewer lysosomes with reduced acidity and proteolytic activity, which is a consequence of impaired microtubule severing and endosomal tubule fission. This aligns with the known role of Spastin in lysosomal function, indicating that the Spast–APP interaction has profound implications for understanding neurodegenerative diseases, particularly AD. The traditional amyloid cascade hypothesis focuses on the toxic effects of extracellular Aβ aggregates and intracellular tau aggregation. However, our findings suggest that the loss of normal APP function in maintaining endolysosomal homeostasis may be equally important in the pathogenesis of AD. This is consistent with the growing recognition that endolysosomal dysfunction is an early and prominent feature of AD. Our findings provide a mechanistic explanation for these observations. If APP function is compromised by genetic mutations, proteolytic processing, or other mechanisms, the resulting deficiency in lysosomal biogenesis directly impairs autophagosome-lysosome fusion and autolysosomal maturation. This creates a vicious cycle in which impaired autophagy leads to the accumulation of damaged organelles and protein aggregates, further compromising cellular function and potentially accelerating neurodegeneration.

Conversely, although we have demonstrated that APP deficiency leads to reduced lysosomal numbers and impaired function, a current limitation is that the molecular mechanisms by which APPL modulates Spastin-Chmp1 dissociation and promotes endosomal fission remain incompletely understood. Without structural or mutagenesis data, we cannot rule out indirect effects (e.g., APPL altering membrane curvature, which then affects Spastin activity). Structural characterization of the APPL-Spastin-Chmp1 complex and mutagenesis of candidate interaction surfaces will be required to define the molecular mechanism. Additionally, the transcriptional and post-transcriptional pathways linking APP deficiency to reduced lysosomal biogenesis require further investigation. Structural characterization of the APP-Spastin interaction would provide critical mechanistic insight. Recent studies have highlighted the importance of transcription factors, such as TFEB and TFE3, in the regulation of lysosomal biogenesis ^4,7^. We found that APPL deletion neither reduced *mitf* (a homolog of TFEB) mRNA levels nor altered Mitf subcellular localization, indicating that APPL regulates lysosome function without affecting lysosomal gene expression. While our findings establish a robust mechanism in phagocytic cells, a major limitation is that we have not validated the APP-Spastin pathway in neurons, where APP is most abundant and where its dysfunction is most relevant to Alzheimer’s disease. Additionally, the use of constitutive App knockout mice may have elicited developmental compensatory mechanisms that could obscure or modify the phenotype, necessitating future studies with conditional, tissue-specific knockouts. While we show reduced lysosome numbers in cortical neurons of *app-/-* mice **(Fig. S11)**, we have not tested whether this affects neuronal efferocytosis by microglia or neuronal autophagy, both of which are implicated in AD pathogenesis. Without neuronal functional assays, the AD relevance of our findings remains inferential. Additionally, while we demonstrate AC degradation defects in *app-/-* BMDMs in vitro, we have not performed tissue-specific genetic rescue or comprehensively quantified efferocytosis in vivo across multiple tissues in the mouse model, which limits our ability to assess physiological relevance. Notably, we examined the lysosome numbers of neurons in the mouse cerebral cortex using TEM, while *app* deletion showed significantly reduced lysosomes, providing preliminary evidence of neuronal lysosomal defects. However, a key limitation of the current study is that we have not tested whether the APP-Spastin pathway operates in neurons during efferocytosis by microglia, nor have we validated these findings in AD models. Future studies are needed to address these questions. Understanding the function of the APP-Spastin interaction in neurons and how its disruption contributes to neurodegeneration is critical for developing targeted therapeutic strategies.

While APP has been extensively studied for its proteolytic processing in AD pathogenesis, our results across three model systems (*Drosophila*, *C. elegans*, and mice) revealed striking functional conservation that transcends species-specific processing pathways. This conservation is particularly notable given that *Drosophila* APPL and *C. elegans* APL-1 lack several mammalian-specific APP processing enzymes and pathways, including β-secretase (BACE1) and the full complement of γ-secretase-mediated amyloidogenic processing ^9^. Despite these differences, all three homologs perform the same essential endolysosomal regulatory function through Spastin interaction. This phylogenetic pattern strongly suggests that endolysosomal regulation represents the ancestral, conserved function of APP, whereas amyloidogenic processing is a derived feature that emerged in the mammalian lineage.

The dominant framework for understanding AD pathogenesis, the amyloid cascade hypothesis, focuses primarily on the toxic gain-of-function effects of Aβ aggregates and hyperphosphorylated tau tangles in AD. Our findings reveal a critical loss-of-function mechanism for APP that could contribute to neurodegeneration, potentially acting in parallel with or exacerbating the toxic gain-of-function effects described in the amyloid cascade hypothesis. This "loss-of-function" hypothesis is consistent with the growing recognition that endolysosomal dysfunction is not merely a consequence of AD but rather an early and prominent pathological feature that may precede and contribute to Aβ and tau pathology ^54,55^. It also provides a mechanistic explanation for why genetic variants that reduce APP expression or function (e.g., the protective A673T mutation) may have complex, context-dependent effects on AD risk ^56,57^. Understanding the balance between APP’s protective non-amyloidogenic functions and its pathogenic amyloidogenic processing will be critical for developing effective therapeutic strategies.

We show that APP’s physiological function extends beyond neuronal contexts to include macrophage-mediated clearance, operating through a Spastin-dependent mechanism that is independent of amyloidogenic processing. We revealed that the non-amyloidogenic function of APP serves as a bridge molecule to moderate the interaction between Spastin and Chmp1 for microtubule severing machinery in endolysosomal budding events, to coordinate endosomal tubule fission and lysosome biogenesis, thereby ensuring efficient degradation of ACs. Our findings have significant implications for understanding Alzheimer’s disease pathogenesis. We propose that AD involves a "double-hit" mechanism: loss of normal APP function (impaired lysosome biogenesis via the APP-Spastin) combined with toxic gain-of-function effects of amyloidogenic APP processing (APP-βCTF accumulation, Aβ toxicity). This model provides a mechanistic explanation for the early and prominent endolysosomal dysfunction observed in AD and suggests that therapeutic strategies should aim to restore APP’s physiological functions while reducing its pathogenic processing. Future studies combining *App-/-* mice with Aβ overexpression models (e.g., 5xFAD x *App-/-*) will be required to test whether loss of APP’s lysosomal function synergizes with Aβ toxicity. Beyond AD, the APP-Spastin axis represents a potentially common regulatory node in diverse pathological contexts characterized by impaired efferocytosis or lysosomal dysfunction, including atherosclerosis, autoimmune disorders, and other neurodegenerative diseases. Interestingly, studies reported that APP expression in the salivary glands of Sjögren’s syndrome patients with xerostomia was lower than in healthy subjects, and a subgroup of rheumatoid arthritis patients exhibited abnormal amyloid precursor protein activation ^58,59^. Therapeutic enhancement of Spastin activity or stabilization of the APP-Spastin interaction may offer novel strategies for restoring tissue homeostasis in these disorders. Future studies in microglial and neuronal models, combined with structural characterization of the APP-Spastin interaction and *in vivo* validation in disease models, will be essential to translate these findings into effective therapies.

## Materials and Methods

### Preparation of S2 cells supernatant for IP LC-MS/MS

S2 cells were incubated with 1 dose of ACs or two doses of ACs for 6 h, centrifuged at 1,000 × g for 3 min, and the supernatant was collected for IP LC MS/MS. Plasmids encoding various tagged proteins were co-transfected into S2 cells as previously described, and immunoprecipitation was performed at 4 °C. Briefly, transfected cells were lysed in a buffer containing 50 mM Tris-Cl (pH 7.4), 1% Triton X-100, 0.15 M NaCl, and 1 mM EDTA supplemented with protease inhibitors, and centrifuged to remove debris. The supernatants were incubated with anti-HA magnetic beads (CST #11846) for 2 h at 4 °C. The beads were then washed thrice with TBS (50 mM Tris, 150 mM NaCl) and eluted by boiling in 2× SDS loading buffer for 5 min. For LC-MS/MS, 5 μL of the eluates were separated by SDS-PAGE and visualized by silver staining. The remaining samples were submitted to HOOGEN BIOTECH (Shanghai, China) for LC-MS/MS analysis. Each LC-MS/MS experiment was performed at least three times independently.

### Cell cultures and transfection

*Drosophila melanogaster* Schneider’s S2 cells were cultured in Sf-900 II SFM medium (Gibco, #10902088) supplemented with 1% penicillin/streptomycin. Cells were maintained at 25 °C and transfected at 60–75% confluency using a DDAB suspension (250 μg/mL) ^60^. Human embryonic kidney HEK 293T cells and mouse macrophage RAW 264.7 cells were cultured in DMEM (Gibco) containing 10% fetal bovine serum (FBS) and 1% penicillin–streptomycin, while Jurkat cells were cultured in RPMI-1640 medium (Gibco) with 10% FBS and 1% penicillin–streptomycin. Mammalian cells were incubated at 37 °C in a 5% CO₂ atmosphere. Transfection was performed using Lipofectamine LTX and Plus Reagent (Invitrogen, #15338100) according to the manufacturer’s instructions.

### Generation of stable cell lines

Stable CharON cell lines were generated as described above, according to a previous method ^35^. Briefly, a caspase-activated GFP (GC3AI) and a pH-insensitive mApple (pHlorina) were cloned into pAC5.1 and pCDNA3.1 vectors, respectively, and expressed in S2 cells and Jurkat cells. After 48 h of expression, a single cell was selected and screened in a 96-well plate to generate the CharON stable cell lines.

### Double-stranded RNA production and RNAi treatment

Primers for double-stranded RNA (dsRNA) synthesis were obtained from the *D. melanogaster* RNAi Screening Center (DRSC) TRiP Functional Genomics Resources (Harvard). edu). A T7 promoter sequence was appended to the 5’-end of each primer. dsRNA was synthesized using 1 µg of PCR product and a High Yield RNA Synthesis Kit (NEB #E2040S). The sequences are listed in Supplementary **Table 3**. S2 cells were treated with 10 µg/mL dsRNA and incubated for 48 h to achieve an efficient knockdown. Cells were harvested for RNA and protein extraction or subjected to efferocytosis assays using a confocal microscope (C2, Nikon Instruments).

### Quantification of tubulin length in S2 cells

Fluorescence images of S2 cells expressing Tubulin-mStaygold and Spastin-mCherry (or Spastin-MIT-mCherry, Katanin-mCherry, and Fidgetin-mCherry) were captured using a 63× objective on a confocal microscope (C2, Nikon Instruments). The mean length of tubulin in a cell was measured by drawing a segmented thin line along the tubules using Fiji software (v.1.54, AnalyzeSkeleton). Approximately 10–20 cells were quantified for each sample.

### Generation of *appl^ko^* flies using CRISPR–Cas9

CRISPR-Cas9–mediated *appl^ko^* flies were generated using two sgRNAs designed to target regions containing NGG PAM sites. The sgRNA sequences were as follows: appl sgRNA 1F: gtcgCTGCCAGCGGGTGCAGCTTC; appl sgRNA 2R: aaacGAAGCTGCACCCGCTGGCAG. sgRNAs were synthesized and annealed as previously described ^61^ and ligated into the pMD18T vector (∼50 ng). The sgRNA cassettes were cloned into an attB vector using the Golden Gate system, verified by sequencing, and injected for transgenesis by Unihuaii Inc. using PhiC31-mediated integration. sgRNA transgenic flies were crossed with the *vasa–Cas9* strain to generate Cas9-edited mutants, which were confirmed by qPCR and sequencing.

### Fluorescent probe for phosphatidylinositol 3,5-bisphosphate in S2 cells

Previous research has shown that PI(3,5)P_2_ was a good probe to label multiple vesicles ^62^. *Dictyostelium* SnxA was a highly selective PI(3,5)P2-binding protein. We cloned the PX sequence from *Dictyostelium* SnxA into the pAC-GFP plasmid to label the cytosolic PI(3,5)P_2_.

### Engulfment and degradation assay for the *S. aureus* strain *RN4220*

For bacterial engulfment and degradation assays, *S. aureus* strain *RN4220* was transformed with the plasmid pRN11, which expresses red fluorescent protein. *S. aureus* cells were harvested with an optical density of 0.4–0.6 at 600 nm and centrifuged at 8,000 × g for 10 min at 20 °C. The cells were washed thrice with deionized water and resuspended in 1 mL of Sf–900TM II SFM. S2 Cells were plated at a density of 1 × 10^6^ cells and incubated for 6 h with 20 μL of the *S. aureus* culture. After this period, the cells were mounted in Vectashield medium (Vector Laboratories) and observed under a confocal microscope (C2, Nikon Instruments) with a 40× objective.

### Protein expression and Western analysis

Cells or tissues were lysed on ice in a buffer containing 50 mM Tris-HCl (pH 7.4), 1% Triton X-100, 150 mM NaCl, and 1 mM EDTA, and supplemented with protease inhibitors. The lysates were centrifuged at 20,000 × g for 30 min, and the supernatants were quantified using the bicinchoninic acid (BCA) assay. Equal protein amounts were resolved by 10–15% SDS-PAGE and transferred to PVDF membranes (Millipore). Membranes were blocked in 5% nonfat milk in TBST and incubated overnight at 4 °C with the following primary antibodies: mouse anti-Actin (CST, 1:1000), anti-Flag (Sigma, 1:1000), anti-HA (CST, 1:1000), anti-Crq (1:500), and anti-APP (1:400). HRP-conjugated secondary antibodies (Jackson ImmunoResearch) were used at a dilution of 1:10 000, and the signals were visualized using ECL reagents (Pierce). All quantitative Western blot assays were independently repeated at least thrice.

### Embryo immunostaining

Immunostaining was performed as previously described ^63,64^. Briefly, stage 13 *Drosophila* embryos were fixed in 4% paraformaldehyde and incubated with primary antibodies: anti-Crq (1:400) to label macrophages and anti-Dcp1 (1:400) to label ACs. Secondary antibodies, including goat anti-rabbit Alexa Fluor Plus 594 or anti-mouse Alexa Fluor Plus 488 (Jackson ImmunoResearch Laboratories), were applied at a 1:1000 dilution. Stained embryos were mounted in VectaShield medium and imaged using a confocal microscope (C2, Nikon Instruments) at 20× and 100× magnifications. Images were processed using Adobe Illustrator. The phagocytosis ratio was calculated as the proportion of engulfed ACs relative to the total number of ACs, and the PI (phagocytosis index) was determined as previously described ^63^. PI values were quantified from confocal image stacks obtained from five embryos, with three image stacks analyzed for each.

### Bimolecular fluorescence complementation assay

BiFC analysis was used to visualize protein–protein interactions in living cells. Two fragments of mVenus are fused to the candidate interacting proteins, forming a fluorescent complex upon interaction ^65^. The corresponding coding sequences were cloned into pcDNA3.1-mVenusN and pcDNA3.1-mVenusC vectors, which express the N-terminal (mVN) and C-terminal (mVC) fragments of mVenus, respectively. The constructs were co-transfected into 293T cells using the polyethyleneimine (PEI) transfection method. After 48 h of culture, the fluorescence signals were examined using confocal laser microscopy. Interacting protein pairs were identified by reconstitution of the yellow fluorescence.

### GST pulldown assay

Recombinant GST-tagged bait proteins and His- or MBP-tagged prey proteins were expressed in *E. coli BL21(DE3)* cells following induction with 0.5 mM IPTG at 16 °C for 16 h. Bacterial pellets were lysed by sonication in lysis buffer (50 mM Tris-HCl pH 7.5, 150 mM NaCl, 1 mM DTT, 1% Triton X-100, and protease inhibitor cocktail). Cleared supernatants were incubated with glutathione–Sepharose 4B beads (GE Healthcare) at 4 °C for 2 h to immobilize the GST fusion proteins. After extensive washing with lysis buffer, the bead-bound GST or GST-bait proteins were incubated with equal amounts (5 µg) of purified prey proteins in binding buffer (20 mM Tris-HCl pH 7.5, 100 mM NaCl, 0.1% NP-40, 1 mM EDTA) at 4 °C overnight with gentle rotation. Beads were then washed five times with binding buffer to remove nonspecifically bound proteins. Retained protein complexes were eluted by boiling in 2 × SDS sample buffer, resolved by SDS-PAGE, and analyzed by immunoblotting using anti-GST and anti-His antibodies. GST alone was used as a negative control to exclude nonspecific binding to the resin, and input lysates (10% of total) were loaded as positive references for prey expression. All pull-down assays were independently repeated at least three times.

### Yeast two-hybrid assay

Binary yeast two-hybrid assays for transmembrane proteins were performed using the split-ubiquitin system, in which genes of interest were inserted into Cub or Nub vectors and transformed into the Saccharomyces cerevisiae strain THY.AP4 ^66,67^. Yeast competent cells were prepared using LiAc solution as previously described, followed by transformation with Cub- or Nub-fused constructs. The transformation mixture was gently mixed and incubated at 30 °C for 30 min, followed by heat shock at 42 °C for 40 min. The yeast cells were then centrifuged at 2,000 × g for 1 min, washed with 100 μL sterile water, resuspended in 50 μL sterile water, plated on SD-Leu/Trp dropout medium, and cultured at 30 °C for 3–4 days. Three representative colonies from each transformation were selected and diluted for spot assays. Protein–protein interactions were identified based on growth on SD-Ade/His/Trp/Leu selective plates.

### Yeast three-hybrid assay

*S. cerevisiae* strain Y2HGold was used for the Y3H assay. cDNA of Spas was fused to the pGADT7 vector. pBridge contains two multiple cloning sites, MCS I and MCS II. Chmp1 cloned in the MCS I is fused with the DNA binding domain of pBridge, and APPL cloned in the MCS II is controlled by a MET17 promoter, while APPL expression is controlled by methionine. Plasmids were co-transformed into yeast via the LiAc/PEG method, and transformants were selected on SD/-Trp/-Leu medium. Positive colonies were normalized to OD_₆₀₀_ = 0.1, serially diluted, and spotted onto SD/-Trp/-Leu/-His plates and SD/-Trp/-Leu/-His/-Met plates, respectively. Growth after 3–5 days at 30 °C indicated ligand-dependent reconstitution of transcriptional activation. Empty pGADT7 plasmid transformants served as negative controls for autoactivation and nonspecific interactions. Three independent transformants were tested in triplicate for each condition.

### CuSO_4_-induced BiFC assay in S2 cells

Culture and transfection of S2 cells were described as mentioned above. We construct pMT-Actin-mVN and pMT-Tsr-mVC (backvector is pMT-puro, Actin and Tsr were used as positive control), pMT-Spas-mVN, pMT-Chmp1-mVC, pMT-Appl-mVN, and pMT-Rab7-mVC. The paired vectors were transfected into S2 cells using the DDAB method, and CuSO_4_ was added to the transfected S2 cells 24 hours later (final concentration was 500 mM). After 24 h, the fluorescence signals were examined using confocal laser microscopy.

### Microscopy and imaging analysis

Confocal microscopy images were captured using an inverted laser scanning confocal microscope (C2, Nikon Instruments) with 488 nm (emission filter BP 503–530 nm) and 543 nm (emission filter BP 560–615 nm) lasers. Images were processed and viewed using a confocal microscope (C2, Nikon Instruments). Images of the S2 cells were captured at room temperature.

### Quantification of protein co-localization

Fluorescence images of S2 cells expressing various vectors were captured using a confocal microscope (C2, Nikon Instruments) at 100× magnification. The images were processed using Adobe Illustrator. 3D reconstruction images were obtained using Volocity software (v.6.3, PerkinElmer). The percentage of lysosomes containing hGal3-mCherry foci was manually quantified within a unit area (100 μm^2^) for each strain. To quantify the co-localization of mCherry-Rab5, mCherry-Rab7, mCherry-Lamp1, and mCherry-VPS4^EtoQ^ with Spastin-GFP, Spastin-mCherry, and Chmp1-GFP, fluorescence images were captured using a confocal microscope (C2, Nikon Instruments) at 100× magnification. Co-localization was measured using Fiji software (v. 1.54 (Pearson’s correlation). Approximately 30 cells were quantified per group.

### Lysosome assays

The following staining assays were performed to examine lysosomal acidity and proteolytic activity in the cells. S2 cells seeded on glass-bottom dishes were incubated with LysoTracker Red (L12492, Invitrogen, 100 nM) or Magic Red (ICT-937, Immunochemistry, 1:250) at 25 °C for 10 min. The cells were washed with medium and live imaged immediately using a confocal microscope (C2, Nikon Instruments) at 100× magnification. Four independent experiments were performed on WT and *appl^ko^* cells.

### Statistical analysis

Statistical analyses were performed using the GraphPad Prism software. Data are presented as the mean ± SEM. One-way analysis of variance (ANOVA) or Student’s two-tailed unpaired t-test was used to determine statistical significance. “*” stands for *p* < 0.05, “**” for *p* < 0.01, “***” for *p* < 0.001, and “****” for *p* < 0.0001. For efferocytosis statistics in fly embryos, at least five embryos from each genotype were tested, unless otherwise indicated.

### Ethics statement

All experiments were performed in accordance with the approved guidelines. The animal experiments were performed in accordance with the National Institutes of Health’s Guide for the Care and Use of Laboratory Animals and were approved by the Animal Care and Use Committee of Shaanxi Normal University (Xi’an, China).

## Acknowledgments

We thank the Bloomington *Drosophila* Stock Center for providing *Drosophila melanogaster* strains. This work was supported by the following funding:

National Natural Science Foundation of China grant 32370799 (XH)

National Natural Science Foundation Key Project of China grant 91954114 (XH)

National Natural Science Foundation of China Youth Program grant 31801164 (ZQ)

Program of Innovative Research Team for the Central Universities grant GK202302003 (XH)

## Author contributions

Conceptualization and experimental designing: XH, ZQ

Experiments performation: ZQ, LFQ, YL, LZL, LHB, CZ, ZQY, YQY

Methodology and materials preparation: MXY, LFQ, YL, WH, YQY

Data analysis and experimental advices: ZQ, LFQ, YL, WH, YQY

Writing – original draft: ZQ, XH

Writing – review & editing: ZQ, XH, WH, YQ

Funding acquisition: XH, ZQ

## Declaration of interests

The authors declare that they have no competing interests.

### Data and materials availability

All data needed to evaluate the conclusions in the paper are present in the paper and/or the Supplementary Materials. The mass spectrometry proteomics data have been deposited to the ProteomeXchange Consortium via the PRIDE ^68^ partner repository with the dataset identifiers PXD078974 and PXD080118. Source data were provided in this paper.

---

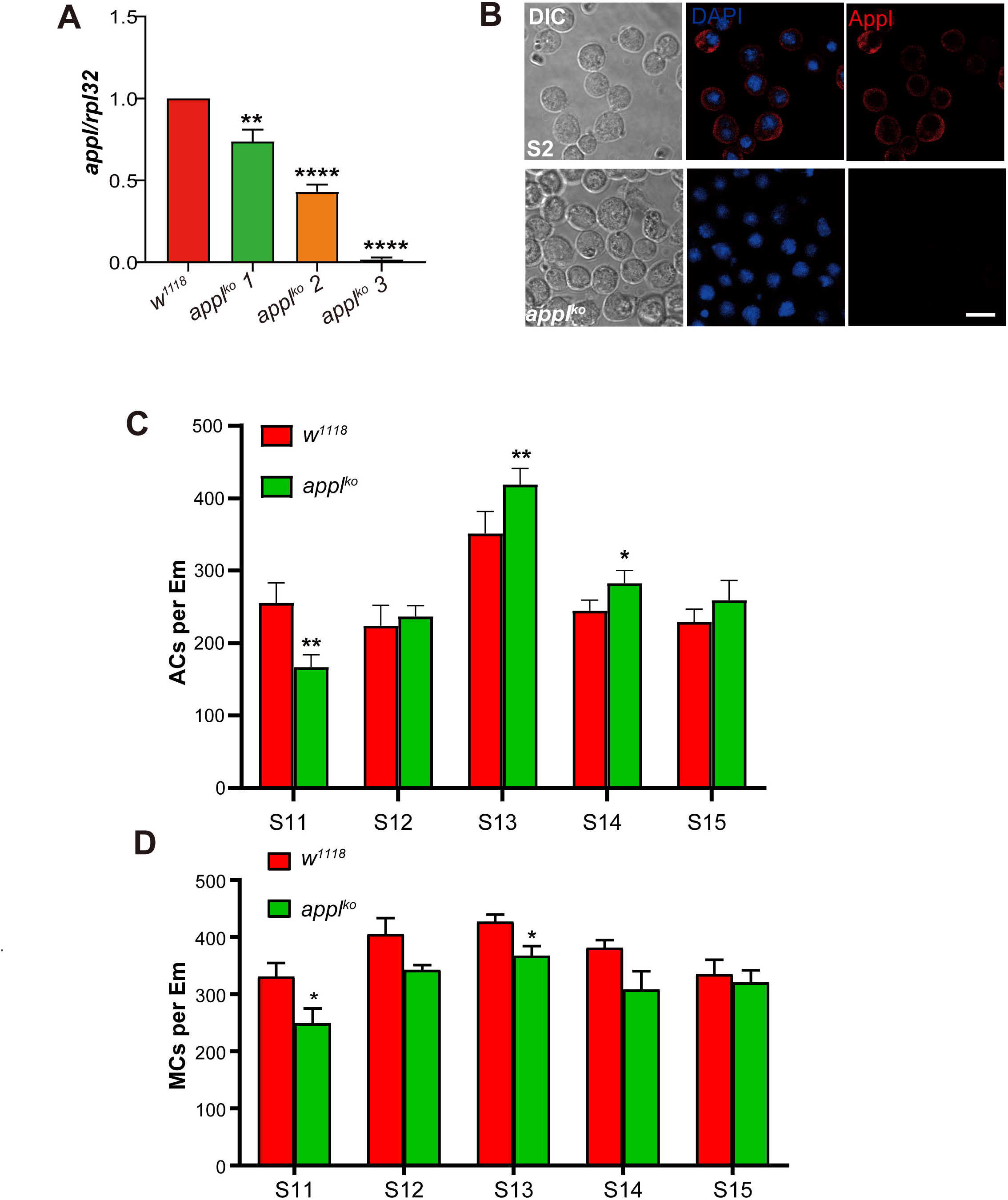

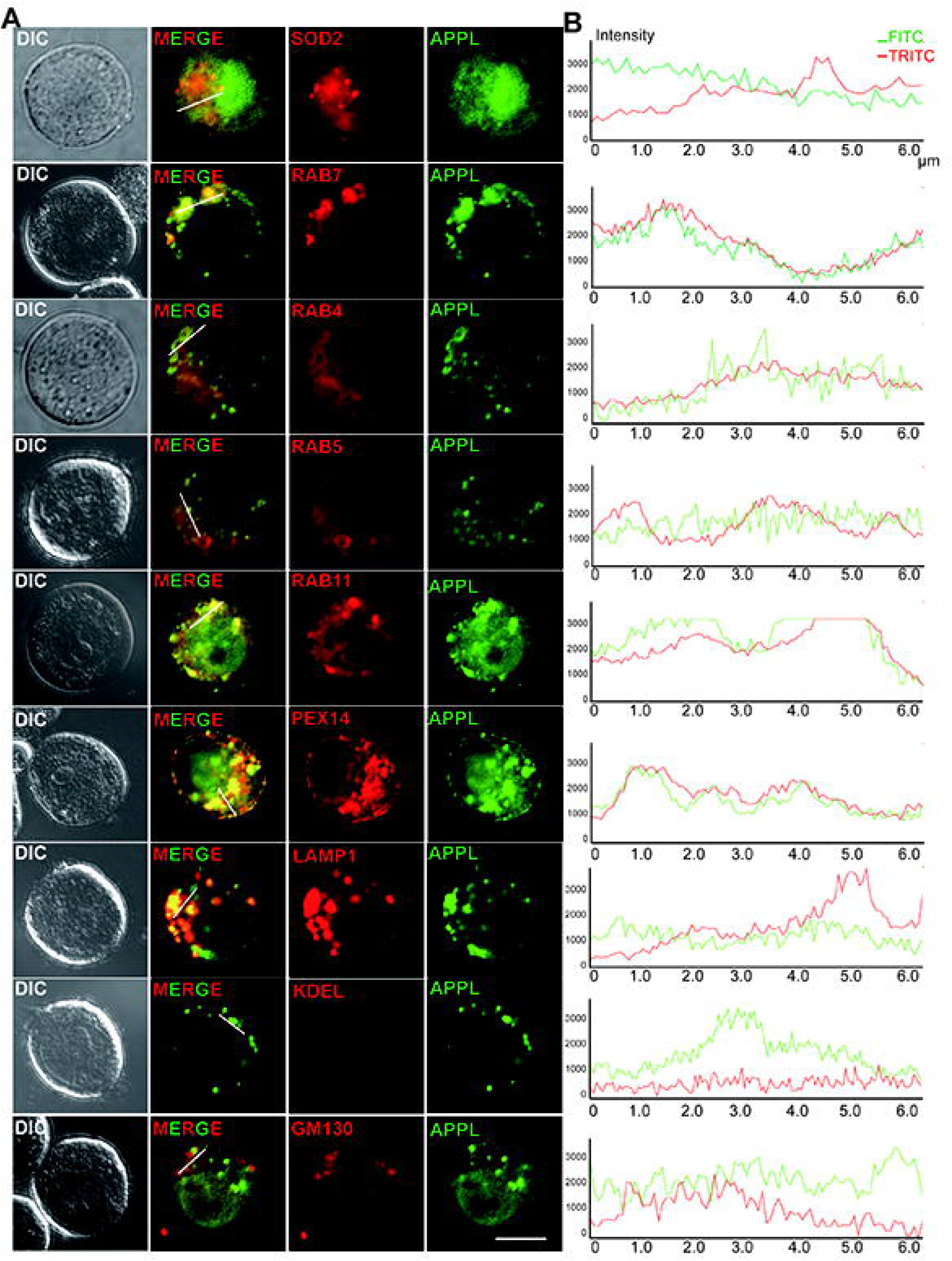

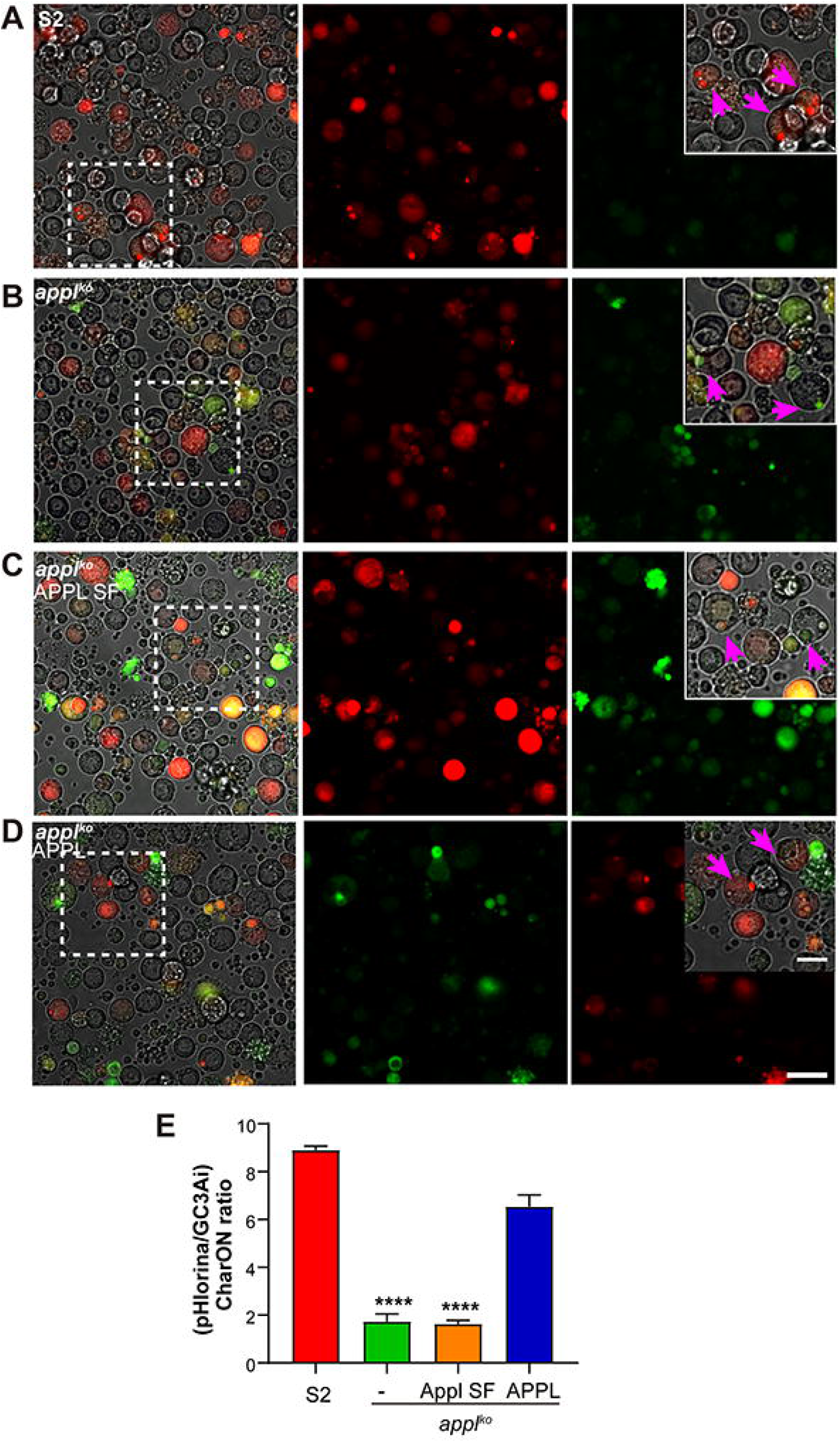

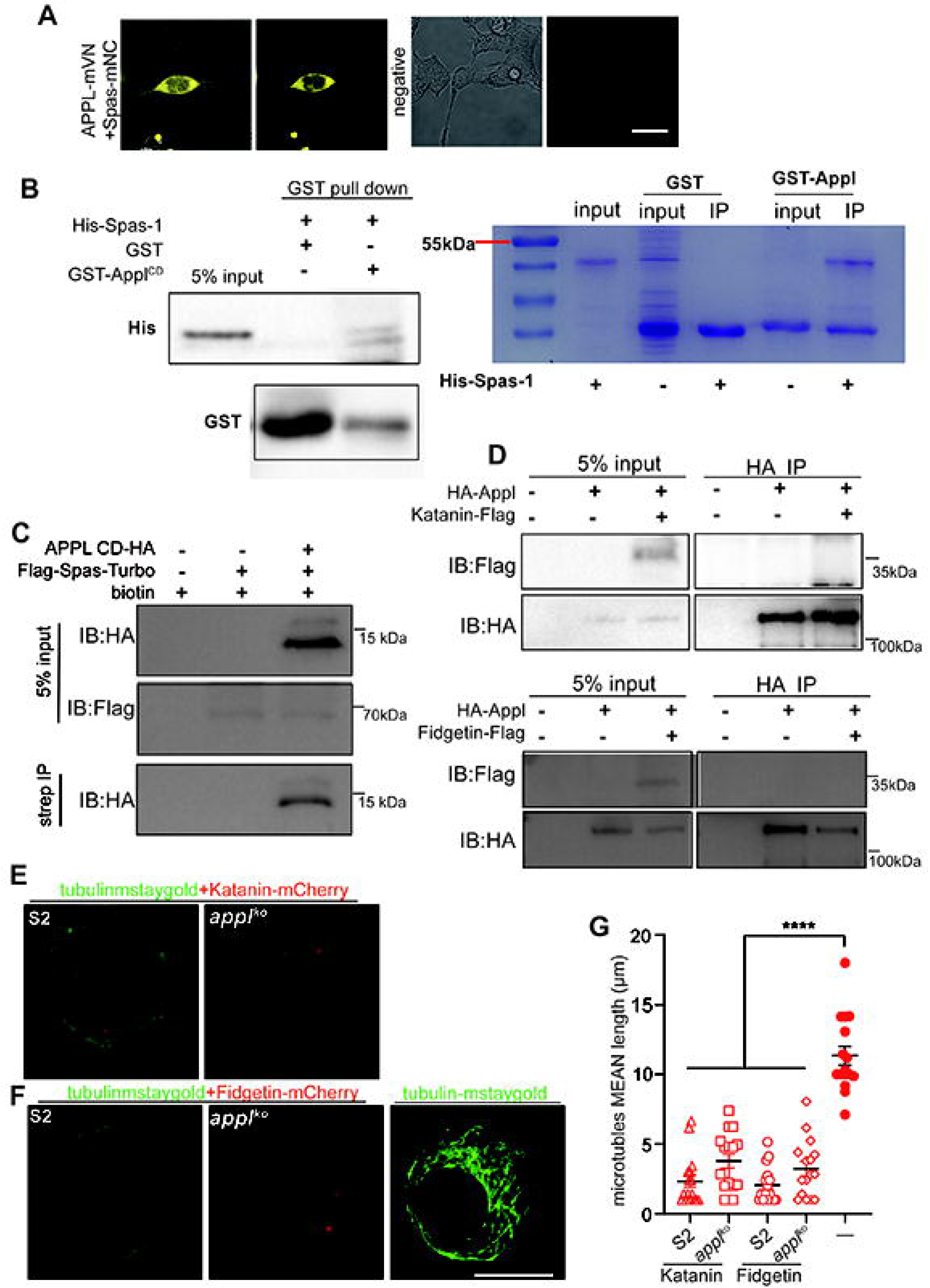

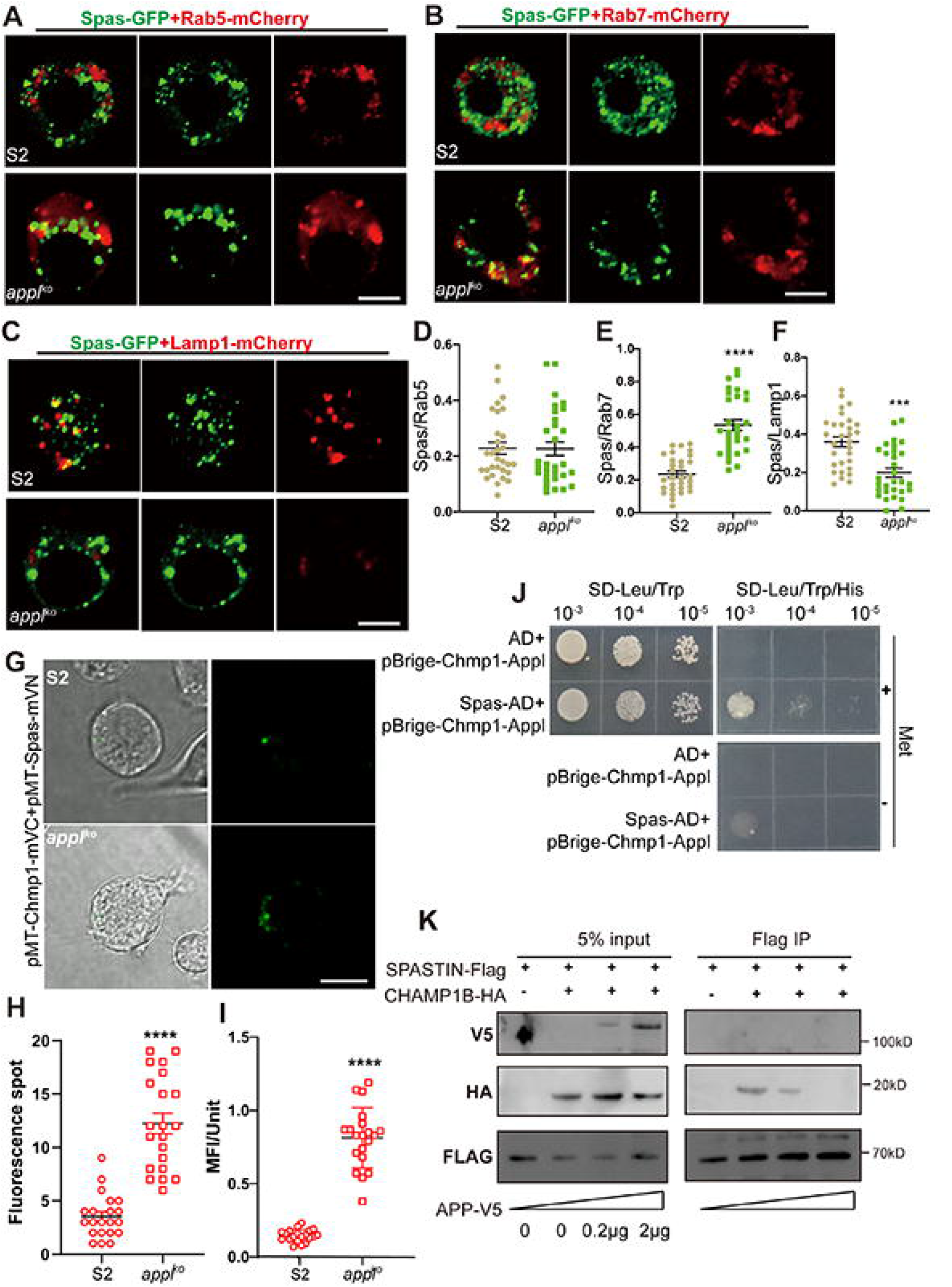

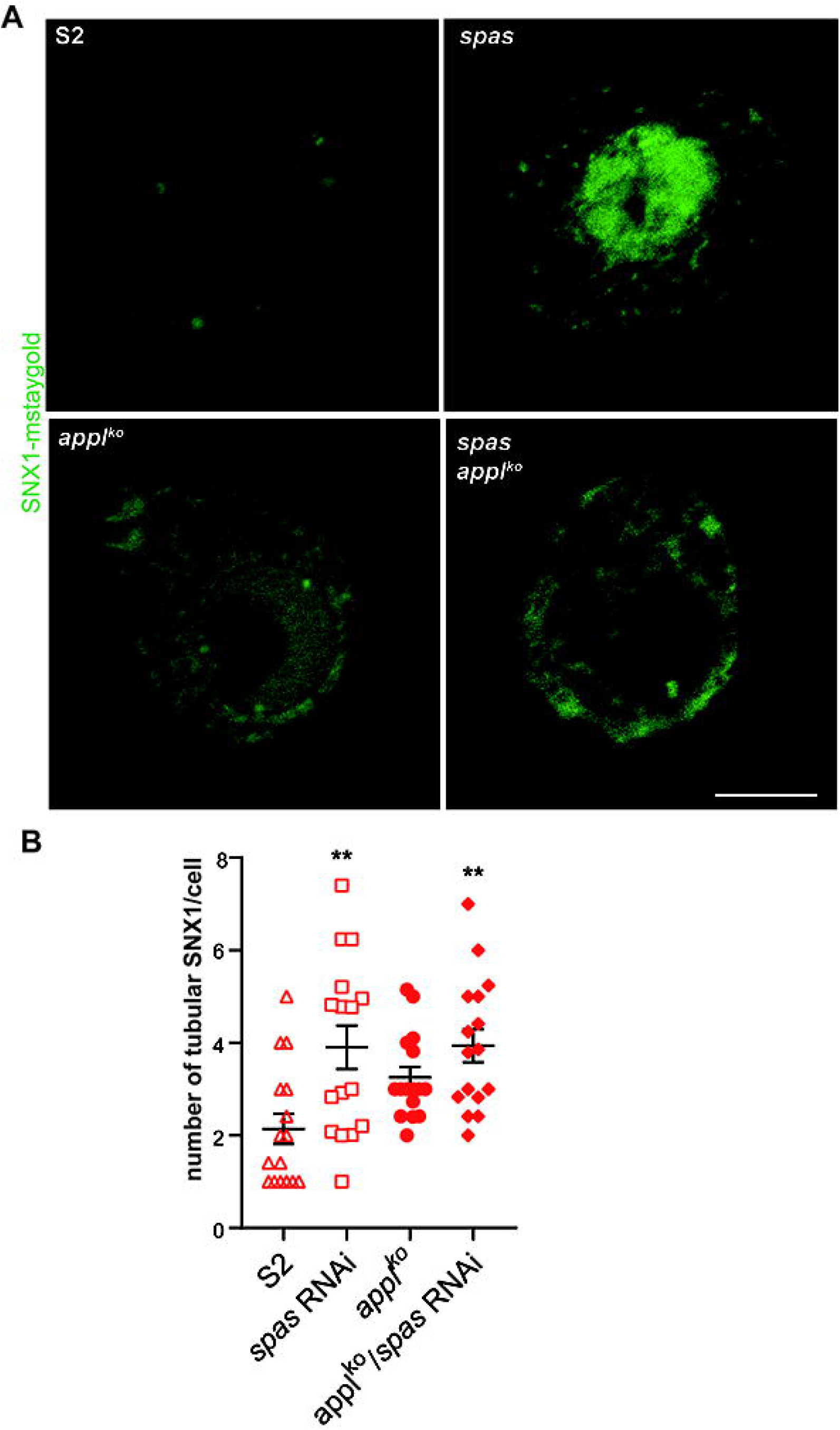

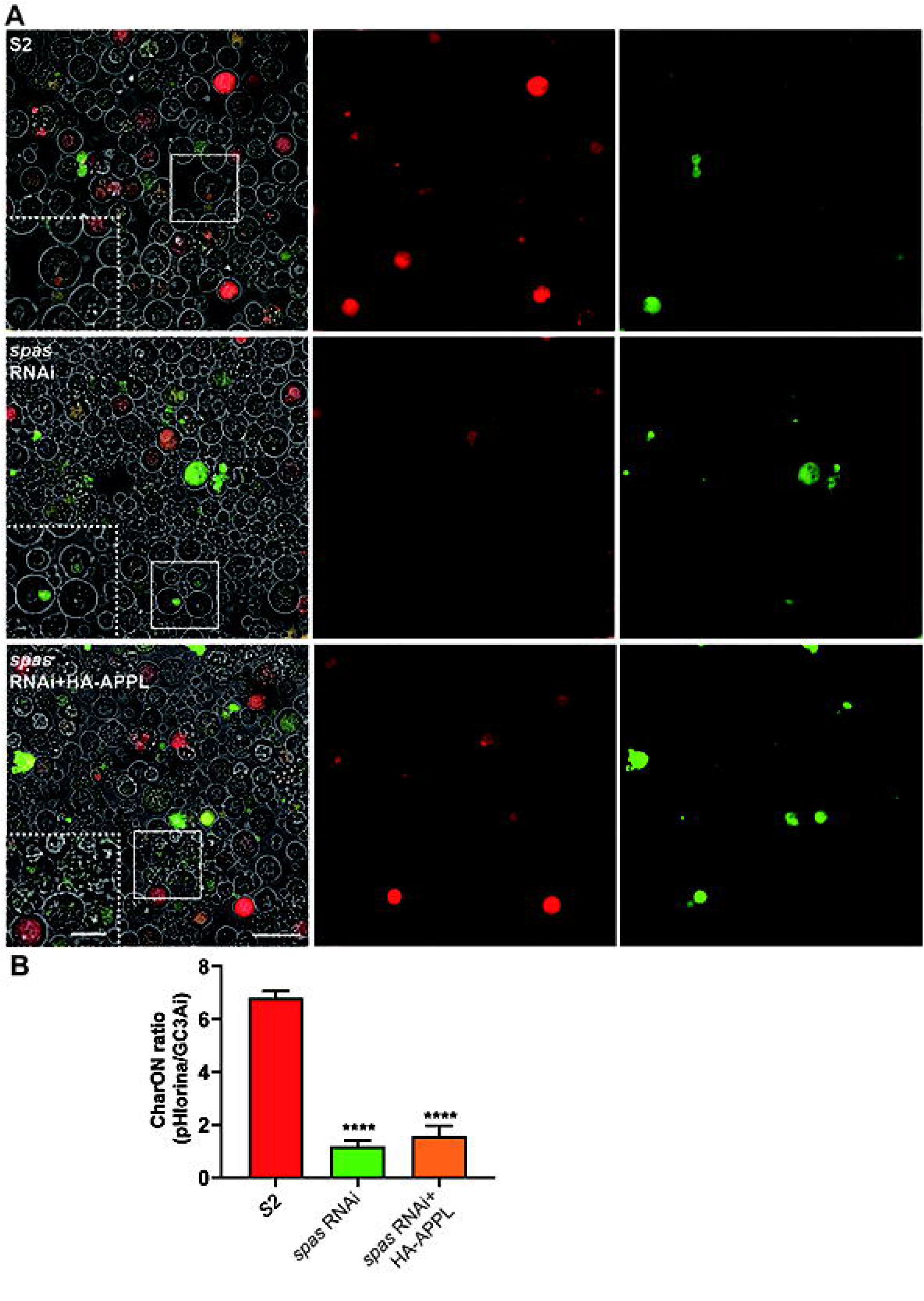

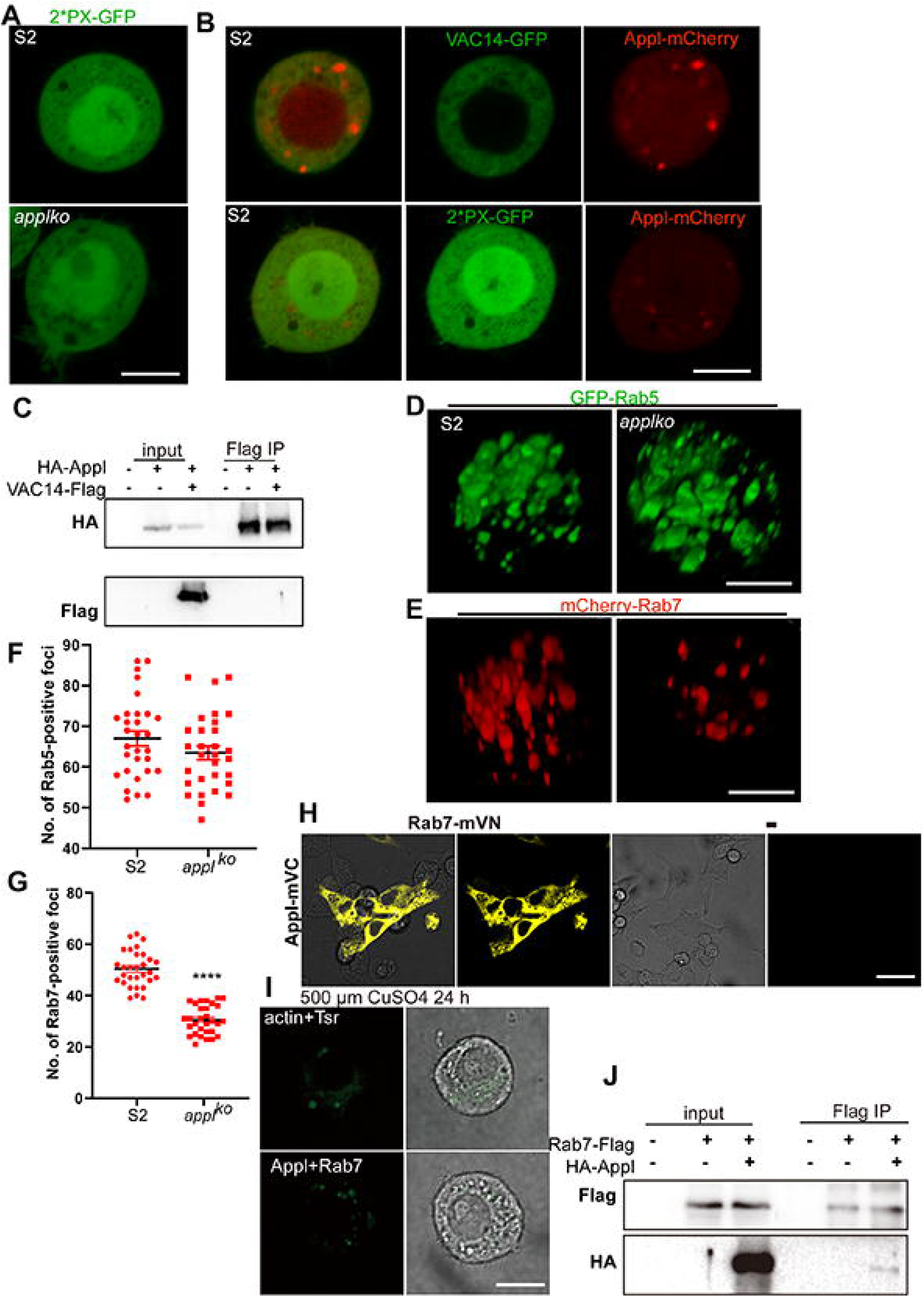

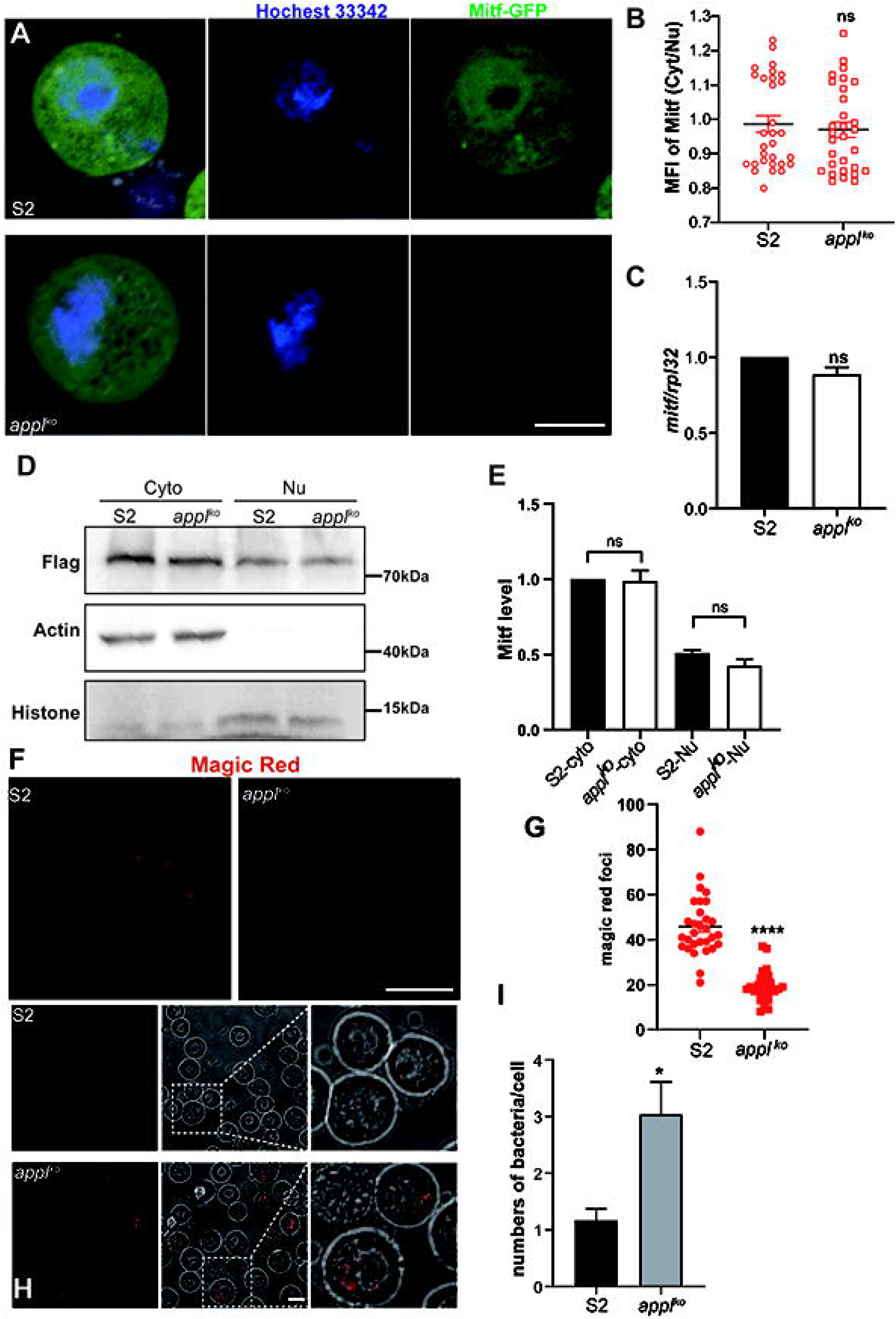

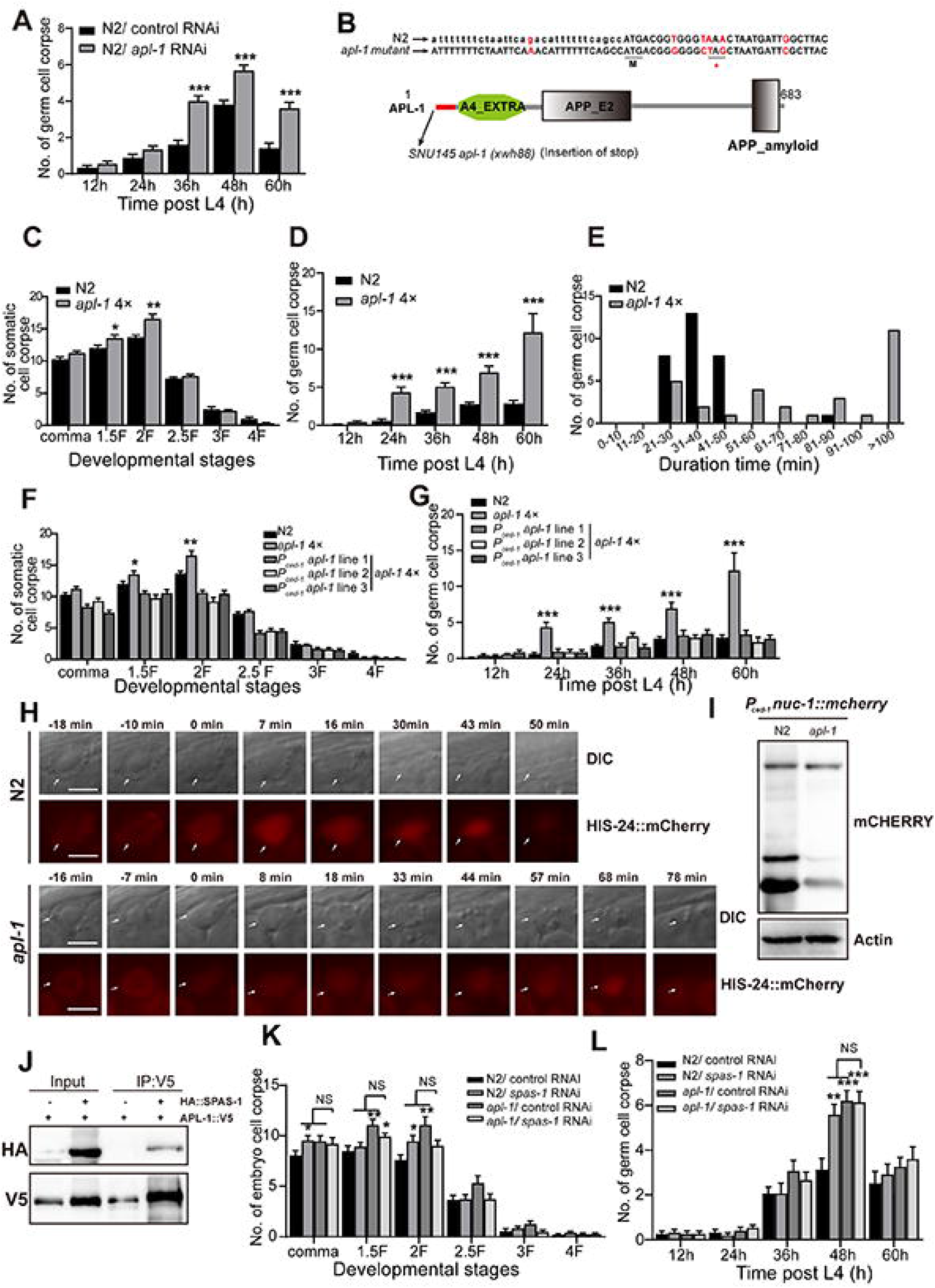

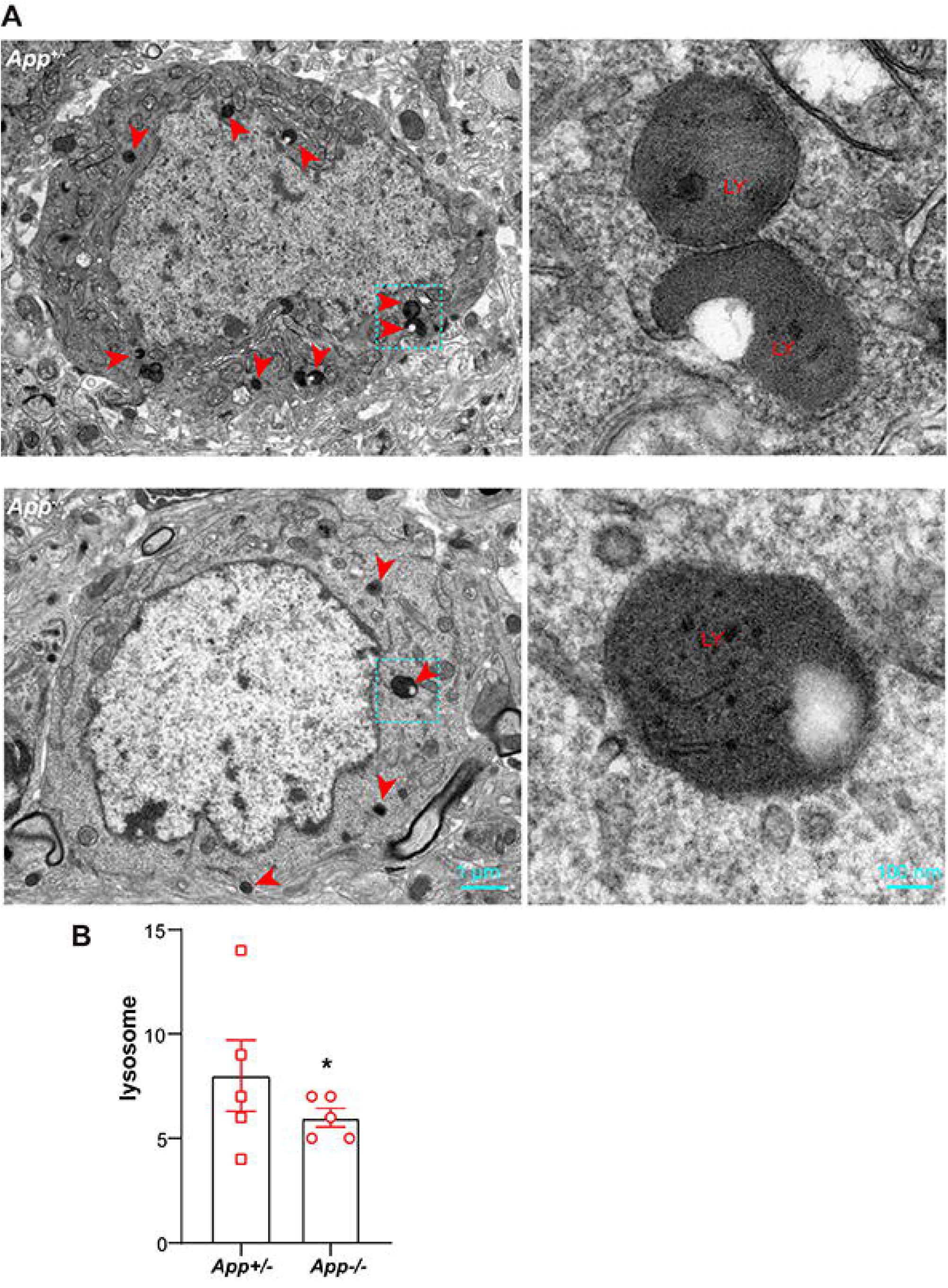

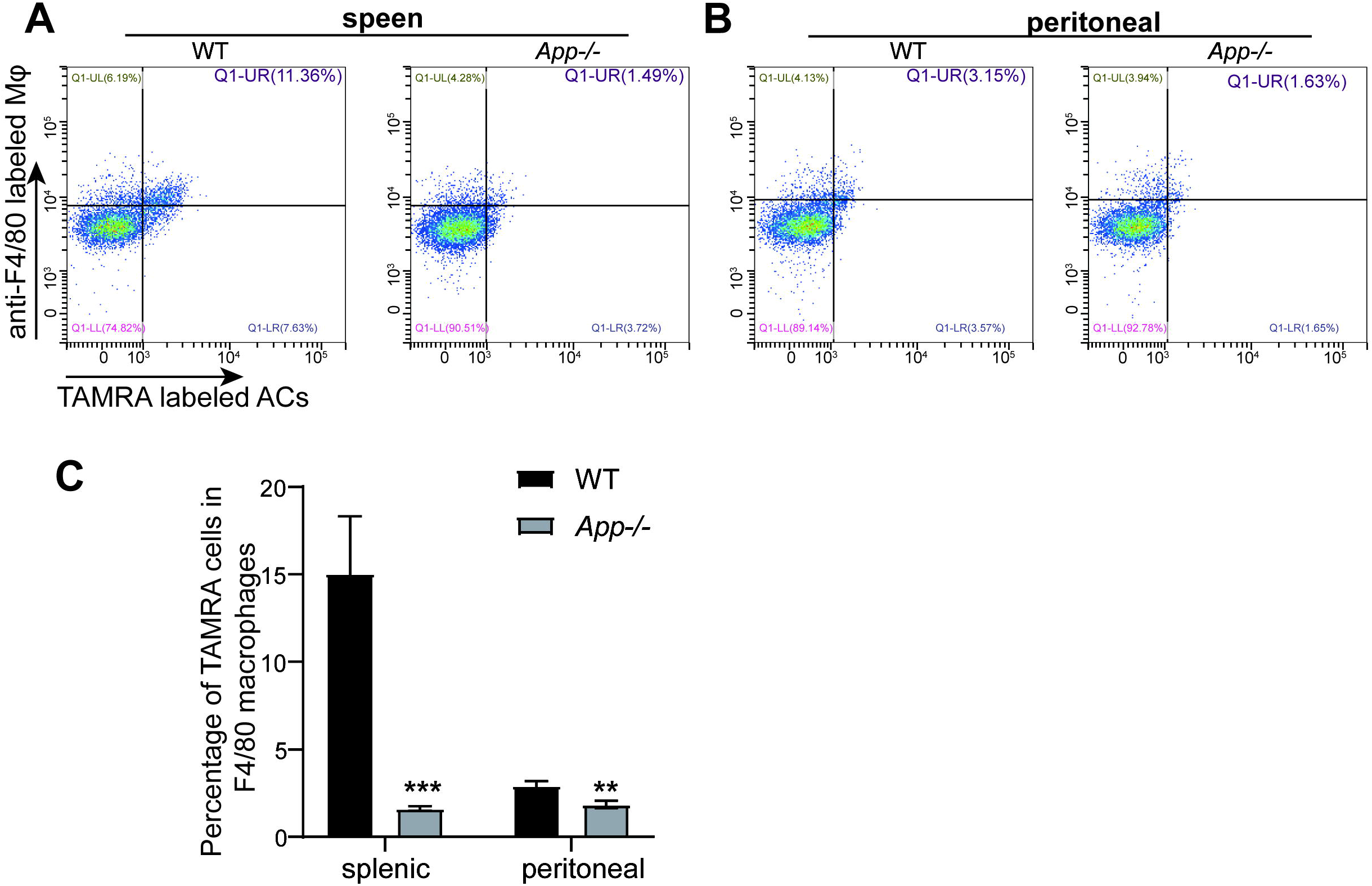

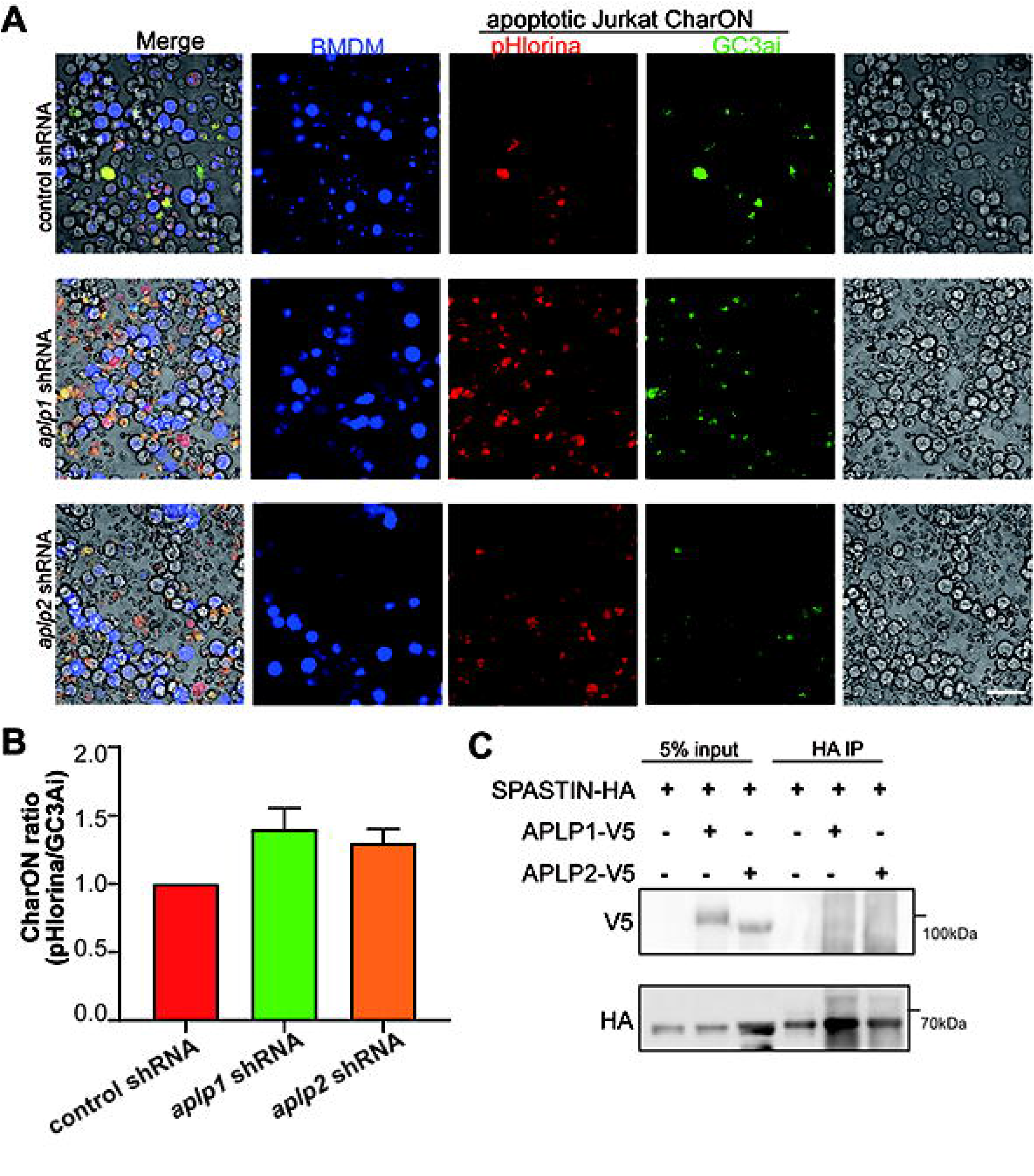

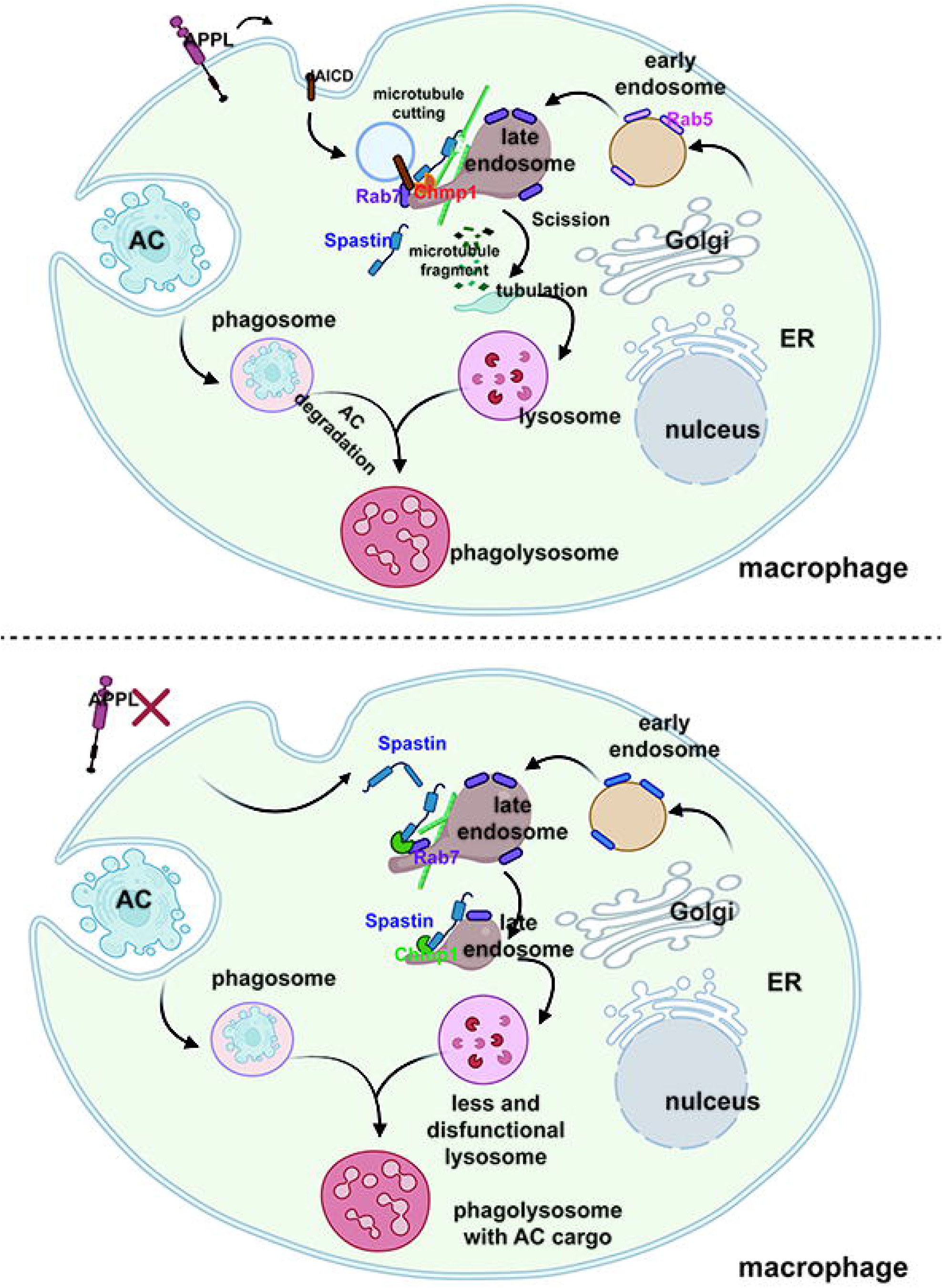

