## Supplemental Table 1-6 for "The evolutionarily conserved APP-Spastin cooperation regulates endolysosomal homeostasis and apoptotic cell degradation": Supplementary files.docx

The **Supplementary Tables file** contain the Supplementary Tables mentioned throughout the article in separated tabs. Below are the corresponding legends.

**Supplementary Table 1. The** **secretory proteins proteomics in phagocytic S2 cells.**

**Supplementary Table 2. Screening for the phagocytosis of the 15 secretory proteins in S2 cells.**

**Supplementary Table 3. The** **APPL-** **immunoprecipitation proteomics in S2 cells.**

**Supplementary Table 4. Screening for the phagocytosis of the 10 intermembrane proteins in S2 cells.**

**Supplementary Table 5. Statistics of cell corpes in *C. elegans*.**

**Supplementary Table 6. The key sources and softwares used in this paper.**

**Fig. S1 APPL mediates AC cargo degradation in *Drosophila* embryos and bacterial degradation.**

**(A)**. *appl* mRNA level was analysed by qRT-PCR, *rpL32* was used as an internal control. Stage 13 embryos of *w^1118^* and *appl^ko^* mRNA were used as samples, and three independent experiments were performed. Data are shown as mean ± SEM. ***p* < 0.01; *****p* < 0.0001 (one-way ANOVA, ‌Dunnett's multiple comparison test‌). **(B)**. Confocal images of S2 and *appl^ko^* S2 cells stained with anti-APPL and Hoechst 33342. Scale bars, 10 μm. **(C).** Column diagram indicates the statistical results for ACs per embryo from stage 11 to stage 15 in *w^1118^* and *appl^ko^* flies. At least three independent experiments were performed, and five embryos were counted for each experiment. Data are shown as mean ± SEM. **p* < 0.05, ***p* < 0.01 (Student’s two-tailed unpaired *t*-test). **(D).** Column diagram indicates the statistical results for macrophages per embryo from stage 11 to stage 15 in *w^1118^* and *appl^ko^*. At least three independent experiments were performed, and five embryos were counted for each experiment. Data are shown as mean ± SEM. **p* < 0.05 (Student’s two-tailed unpaired *t*-test).

**Fig. S2 Intracellular APPL co-localizes with various endosomal and lysosomal markers in S2 cells, indicating its presence in internal membrane compartments.**

**(A).** Co-localization of mCherry-tagged Sod (mitochondria) and GFP-tagged APPL in S2 cells. **(B)**. Co-localization of mCherry-tagged Rab7 (late endosome) and GFP-tagged APPL in S2 cells. **(C).** Co-localization of mCherry-tagged Rab4(fast-recycling endosome) and GFP-tagged APPL in S2 cells. **(D).** Co-localization of mCherry-tagged Rab5 (early endosomes) and GFP-tagged APPL in S2 cells. **(E).** Co-localization of mCherry-tagged Rab11 (recycling endosome) and GFP-tagged APPL in S2 cells. **(F).** Co-localization of mCherry-tagged Pex14 (peroxisome) and GFP-tagged APPL in S2 cells. **(G).** Co-localization of mCherry-tagged Lamp1 (lysosome) and GFP-tagged APPL in S2 cells. **(H).** Co-localization of mCherry-tagged KDEL (ER) and GFP-tagged APPL in S2 cells. **(I).** Co-localization of mCherry-tagged GM130 (Golgi) and GFP-tagged APPL in S2 cells. Scale bars, 5 μm.

**Fig. S3 The secreted form of APPL cannot rescue *appl^ko^* defect in AC cargo degradation.**

**(A-D).** Degradation assay in S2, *appl^ko^*, *appl^ko^/*APPL^SF^ (secretion form) and *appl^ko^/*dAICD (the intracellular domain of APPL) cells. The CharON stable S2 cell line, which expressed a caspase-activated GFP and a pH-insensitive mApple (pHlorina), was treated with 0.25 μg/mL actinomycin D for 6 h, and then added to S2, *appl^ko^*, *appl^ko^*(expressing APPL^SF^), *appl^ko^*(expressing dAICD) cells, respectively, and observed by confocal fluorescence microscopy. Scale bars: main fields, 20 μm; insets, 10 μm. The magenta arrows showed engulfed ACs. **(E).** The column diagram shows ratiometric CharON (pHlorina/GC3ai) signal during macrophage engulfment. Three independent experiments were performed. Data are shown as mean ± SEM. *****p* < 0.0001 (one-way ANOVA, ‌Dunnett's multiple comparison test‌).

**Fig. S4 APPL regulates microtubule cutting without microtubule severing enzymes – Katanin and Fidgetin.**

**(A).** TurboID assay showing Spastin interaction with APPL intercellular domain. HA-APPL intercellular domain and Flag-Spastin-TurboID were co-transfected into S2 cells, cells were treated with 50 μM biotin for 10 min before harvest, cell lysates were immunoprecipitated using streptavidin magnetic beads, and analyzed using western blot with anti-HA. **(B).** Co-IP showed that katanin did not interact with APPL. Katanin-Flag and HA-APPL were co-transfected into S2 cells, cell lysates were immunoprecipitated by anti-HA magnetic beads, and analysed using western blot by anti-Flag and anti-HA. **(C).** Co-IP showed that fidgetin did not interact with APPL. Fidgetin-Flag and HA-APPL were co-transfected into S2 cells, and the cell lysates were immunoprecipitated using anti-HA magnetic beads and analyzed using western blotting with anti-Flag and anti-HA antibodies. **(D-E).** Tubulin with C-terminal mStaygold and Katanin or Fidgetin with C-terminal mCherry were co-expressed in S2 and *appl^ko^* cells, respectively. Scale bars, 5 μm. **(F).** The mean length of microtubules per cell was measured in 16 cells; data are shown as mean ± SEM; *****p* < 0.0001 (one-way ANOVA, ‌Dunnett's multiple comparison test‌).

**Fig. S5 The interaction of Spastin and Chmp1 was enhanced by the absence of APPL.**

**(A)**. Co-localization of Spastin-GFP and Rab5-mCherry in S2 and *appl^ko^* cells. Scale bars, 5 μm. **(B)**. Co-localization of Spastin-GFP and Rab7-mCherry in S2 and *appl^ko^* cells. Scale bars, 5 μm. **(C)**. Co-localization of Spastin-GFP and Lamp1-mCherry in S2 and *appl^ko^* cells. Scale bars, 5 μm. **(D)**. The extent of co-localization between Spastin and Rab5 proteins was estimated by calculating the Pearson’s correlation coefficient for green and red pixels in each cell, using Fiji software (n = 30 cells, *n* = 10 cells in each independent experiment), data show mean ± SEM (Student’s two-tailed unpaired t-test). **(E)**. The extent of co-localization between Spastin and Rab7 proteins was estimated by calculating the Pearson’s correlation coefficient for green and red pixels in each cell using Fiji software (n = 30 cells, *n* = 10 cells in each independent experiment); data are shown as mean ± SEM, *****p* < 0.0001 (Student’s two-tailed unpaired t-test). **(F)**. The extent of co-localization between Spastin and Lamp1 proteins was estimated by calculating the Pearson’s correlation coefficient for green and red pixels in each cell using Fiji software (n = 30 cells, *n* = 10 cells in each independent experiment); data are shown as mean ± SEM, ****p* < 0.001 (Student’s two-tailed unpaired t-test). **(G)**. CuSO_4_ induced BiFC showed Spastin and Chmp1 in S2 and *appl^ko^* cells. pMT-Spastin-mVN and pMT-Chmp1-mVC were co-transfected into S2 cells, 500mM CuSO_4_ was added to allow for 24 h incubation, and mVenus fluorescence images were captured by confocal microscope. Scale bars, 10 μm. **(H and I).** The number of mVenus foci per cell and MFI per cell is shown in the column, n = 15 cells in each of the three independent experiments, data are shown as mean ± SEM, *****p* < 0.0001 (Student’s two-tailed unpaired *t*-test). **(J)**. Yeast three-hybrid assays were used to detect interactions between APPL, Chmp1 and Spastin transformants. Different concentrations of labeled yeast transformants were assayed on SD- Leu/Trp/His and SD-Leu/Trp/His/Met plates for growth. Empty AD and pBridge-APPL-Chmp1 were used to detect self-activation of the probe. **(K).** Protein interaction competition experiment showed that APP inhibited SPASTIN and CHMP1B interaction. SPASTIN-Flag, APP-V5 and CHMP1B-HA were expressed and purified from 293T cells, 1 μg SPASTIN and 2 μg CHMP1B was quantified to detect the interaction, after 1 h incubation, 0 ng, 200 ng and 2 μg purified APP was added into the mixture, respectively, and the protein mixture was immunoprecipitated with anti-Flag magnetic beads and analyzed using western blotting with anti-Flag, anti-V5 and anti-HA antibodies.

**Fig. S6 Spastin and APPL regulates endosomal tubulation.**

**(A).** Transfect SNX1-mstaygold into S2, *spastin* RNAi, *appl^ko^*, and *appl^ko^/spastin* cells. Scale bars: main fields, 5 μm; insets, 2 μm. **(B).** The mean number of SNX1 tubules per cell was quantified (*n* = 15 cells counted per experimental condition in each experiment). Data are shown as mean ± SEM, ***p* < 0.001 (one-way ANOVA, ‌Dunnett's multiple comparison test‌).

**Fig. S7 APPL cannot rescue the defect in AC cargo degradation caused by *spastin* knockdown.**

**(A).** Degradation assay in S2, *spastin* RNAi, and *spastin* RNAi/HA-APPL cells. The CharON stable S2 cell line, which expressed a caspase-activated GFP and a pH-insensitive mApple (pHlorina), was treated with 0.25 μg/mL actinomycin D for 6 h, and then added to S2, *spastin* RNAi, and *spastin* RNAi/HA-APPL cells, respectively, and observed by confocal fluorescence microscopy. Scale bars: main fields, 20 μm; insets, 10 μm. The magenta arrows showed engulfed ACs. **(B).** The column diagram shows ratiometric CharON (pHlorina/GC3ai) signal during macrophage engulfment. Three independent experiments were performed. Data are shown as mean ± SEM. *****p* < 0.0001 (one-way ANOVA, ‌Dunnett's multiple comparison test‌).

**Fig. S8 APPL interacts with Rab7 and regulates lysosomal numbers.**

**(A)**. Localization of 2*PX-GFP in S2 and *appl^ko^* cells. Scale bars, 5 μm.  **(B)**. Co-localization of vac14-GFP, 2*PX-GFP and APPL-mCherry in S2 cells. Scale bars, 5 μm.  **(C)**. Co-IP showed that APPL did not interact with VAC14 in S2 cells. APPL-HA and VAC14-Flag were co-transfected into S2 cells, and the cell lysates were immunoprecipitated with anti-HA magnetic beads and analyzed using western blotting with anti-Flag and anti-HA antibodies. **(D)**. Confocal fluorescence images of S2 and *appl^ko^* cells transfected with GFP-Rab5 in the z-axis direction. Scale bars, 5 μm. **(E)**. Confocal fluorescence images of S2 and *appl^ko^* cells transfected with mCherry-Rab7 in the z-axis direction. Scale bars, 5 μm. **(F)**. The number of early endosomes per cell as determined by GFP signals foci is shown, n = 30 cells, data are shown as mean ± SEM, Student’s two-tailed unpaired t-test. **(G)**. The number of early endosomes per cell as determined by mCherry signals foci is shown, n = 30 cells, data are shown as mean ± SEM, *****p* < 0.0001 (Student’s two-tailed unpaired t-test). **(H)**. BiFC showed APPL binding to Rab7 in 293T cells. APPL-mVN and Rab7- mVC were co-transfected into 293T cells, and mVenus fluorescence images were captured by confocal microscope. Scale bars, 20 μm. (I). CuSO_4_ induced BiFC showed that APPL interacted with Rab7 in S2 cells. pMT-APPL-mVN and pMT-Rab7- mVC were co-transfected into S2 cells, 500mM CuSO_4_ was added to allow for 24 h incubation, and mVenus fluorescence images were captured by confocal microscope. Scale bars, 10 μm. **(J)**. Co-IP showed that APPL interacted with Rab7 in S2 cells. APPL-HA and Rab7-Flag were co-transfected into S2 cells, and the cell lysates were immunoprecipitated with anti-HA magnetic beads and analyzed using western blotting with anti-Flag and anti-HA antibodies.

**Fig. S9 APPL regulates lysosomal function in a Mitf-independent manner.**

**(A).** Mitf-GFP was transfected into S2 and *appl^ko^* cells, and the cells were stained with Hochest 33342 for 10 min. Scale bars, 5 μm. **(B).** The ratio of mean fluorescence intensity (MFI) (Cyt/Nu) of Mitf-GFP in S2 and *appl^ko^* cells, *n* = 30 cells, data are shown as mean ± SEM (Student’s two-tailed unpaired *t*-test). **(C).** Column diagram shf Fig. 1F owing the mRNA levels of *mitf* in S2 and *appl^ko^* cells. Three independent experiments were performed, and the data are shown as mean ± SEM, Student’s two-tailed unpaired *t*-test. **(D).** WB showed that Mitf level and location were not affected by APPL. cell lysates were collected from S2 cells and *appl^ko^* cells, and the cell lysates were analyzed using western blotting with anti-Flag, anti-Actin and anti-Histone antibodies. **(E).** Column diagram showing the protein levels of Mitf in S2 and *appl^ko^* cells. Three independent experiments were performed, and the data are shown as mean ± SEM, Student’s two-tailed unpaired *t*-test. **(F).** Confocal fluorescence images of S2 and *appl^ko^* cells stained with Magic Red. Scale bars, 5 μm. **(G).** Quantification of the number of lysosomes stained with Magic Red. *n* = 10 cells in each of the three independent experiments; data are shown as mean ± SEM, *****p* < 0.0001. (Student’s two-tailed unpaired *t*-test). **(H).** *S. aureus* strain RN4220 was added to S2 and *appl^ko^* cells at a ratio of 10:1 to allow phagocytosis for 6 h, then wash out the *S. aureus* was washed out to continue the degradation process for 3h, number of bacteria is shown in **(I).** Scale bars: main fields, 10 μm; insets, 5 µm. Three independent experiments were conducted. Data are shown as mean ± SEM. **p* < 0.05(Student’s two-tailed unpaired *t*-test).

**Fig. S10 APL-1 and SPAS-1 participates in AC degradation in *C. elegans***

**(A).** Quantification (mean ± SEM) of button-like cell corpses at different stages post L4 in N2/control and *apl-1* RNAi worms. Fifteen adult worm germlines were scored at each stage for every strain. **(B).** Sequencing results for the *apl-1* mutant, with the mutated base indicated in red. Schematic illustration of the *apl-1(xwh143)* homozygous mutations generated by CRISPR-Cas9 editing of the endogenous *apl-1* loci. Amino acids near the mutated sites are indicated by arrows. **(C, D, and E).** The development of embryo stages and adult corpses was quantified in the indicated strains. Fifteen adult worms or embryos were scored at each stage for every strain. **(F).** Extrachromosomal expression of *apl-1* driven by the *ced-1* promoter (*P_ced-1_apl-1*, overexpression of APL-1 in phagocytes) resulted in the complete rescue of cell corpse clearance in *apl-1(xwh143)* worms. **(G)** Four-dimensional microscopy analysis of the persistence of 30 germ cell corpses was performed in N2, *apl-1(xwh143)* mutant, and three *P_ced-1_apl-1 lines in apl-1(xwh143)* mutant. **(H).** Time-lapse chasing of HIS-24:: mCherry-positive phagolysosomes in N2 and *apl-1(xwh143)* worm germlines. The time point at which HIS-24::mCherry mostly rings the cell corpse was set to 0 min. Arrows indicate the continuous presence of cell corpses (DIC, top row) and HIS-24::mCherry (fluorescence, bottom row) in the pharynx in the pharynx of the hermaphrodite. DIC and fluorescence images were obtained at specific time intervals. Bars, 5 µm. **(I).** Exogenous NUC-1::mCHERRY protein levels were examined by immunoblot analysis in N2 and *apl-1(xwh143)* whole worm lysates. **(J).** Co-IP showed that APL-1 interacted with SPAS-1 in 293T cells. Apl-1-V5 and HA-SPAS-1 were co-transfected into 293T cells, and the cell lysates were immunoprecipitated with anti-HA magnetic beads and analyzed using western blotting with anti-V5 and anti-HA antibodies. **(K and L).** Quantification of button-like cell corpses at different stages post L4 in N2/control, *spas-1* RNAi, *apl-1(xwh143), and spas-1* RNAi in *apl-1(xwh143)* mutants. Fifteen adult worm germlines were scored at each stage for each strain. Data are shown as mean ± SEM. ns, no significance, **p* < 0.05, ***p* < 0.01, ****p* < 0.001(Student’s two-tailed unpaired *t*-test).

**Fig. S11 The mouse *App-/-* shows significant reduced lysosomes in neurons by TEM.**

**(A).** TEM images of neurons from *App+/-* and *App-/-* mouse cerebral cortex respectively, the red arrows represented lysosomes, the blue dotted-line rectangle was zoomed in the right pannel. Scale bars: main fields, 1 μm; insets, 100 nm. **(B).** The column showed the number of lysosomes per cell as determined by TEM, n = 5 cells, data are shown as mean ± SEM; **p* < 0.05 (Student’s two-tailed unpaired t-test).

**Fig. S12 *In vivo* efferocytosis assay for measuring ACs degradation in tissue-resident macrophages**

**(A and B).** Representative flow cytometry dot plots showing F4/80 expression and TAMRA signal in macrophages from the spleen and peritoneal cavity of wild-type (WT) and *App-/-* mice. TAMRA-labeled apoptotic cells were delivered into mice, and after a standard incubation period, splenic single-cell suspensions and peritoneal lavage cells were collected, stained with anti-F4/80 antibody, and subjected to flow cytometric analysis. The TAMRA signal is enhanced upon lysosomal degradation of phagocytosed apoptotic cells, and therefore reports intracellular degradative efficiency rather than initial engulfment. **(C).** Quantitative summary of the percentage of TAMRA+ cells among *App-/-* macrophages. Data are presented as mean ± SEM, with n = 3 mice per group. Statistical comparisons between genotypes were performed using unpaired two-tailed Student’ s *t*-test for each tissue independently.

**Fig. S13 The two APP-like proteins had no function in AC degradation.**

**(A).** Roles of Aplp1 and Aplp2 in the degradation of ACs by BMDMs. The CharON stable Jurkat cell line, was treated with 10 ng/mL staurosporine for 12 h, and then added to control- *aplp1-* and *aplp2-* RNAi treatment BMDMs, respectively. BMDMs nuclei were stained with DAPI, and observed by confocal fluorescence microscopy. Scale bars, 50 μm. **(B).** Three independent experiments were performed. Data are shown as mean ± SEM, ns: no significance (one-way ANOVA, ‌Dunnett's multiple comparison test‌). **(C)**. Co-IP analysis showed that Aplp1 and Aplp2 did not interact with Spastin. Aplp1-V5, Spastin-HA, Aplp2, and Spastin-HA were co-transfected into 293T cells, and the cell lysates were immunoprecipitated using anti-HA magnetic beads and analyzed using western blotting with anti-V5 and anti-HA antibodies.

**Fig. S14 The model of lysosome function regulated by APP and Spastin.**

**(A).** The microtubule severing protein Spastin binds to the cytoplasmic domain of APP via its MIT domain, forming a scaffold which is recruited to the late endosome membrane via the interaction between APPL and Rab7. Spastin cuts the microtubules surrounding the endosome tubules, the following microtubule cutting promotes endosome scission; Endosomal tubule fission, which may carry essential lysosomal enzymes or receptors is important for the downstream lysosome biogenesis. **(B).** When APP is lost or abnormal, the dissociation of Chmp1 and Spastin is disrupted, which fails to cut microtubule, then increases endosomal tubulation, mistrafficking of cargoes, and dysregulation of proper lysosomal functions, finally resulting in defect of cargo degradation.
